# Phylogenetics and phylogeography of *Euphorbia canariensis* reveal an extreme Canarian-Asian disjunction and limited inter-island colonization

**DOI:** 10.1101/2023.01.27.525850

**Authors:** Alberto J. Coello, Pablo Vargas, Emilio Cano, Ricarda Riina, Mario Fernández-Mazuecos

## Abstract

*Euphorbia canariensis* is an iconic endemic species of the Canary Islands and one of the most characteristic species of lowland xerophytic communities known, in Spanish, as ‘cardonal-tabaibal’. This species is widely distributed in the archipelago, which contrasts with the theoretically low dispersal abilities suggested by its unspecialized diasporas. Furthermore, the phylogenetic relationships of this species are unclear, although it is thought to be related to the Indian *E. epiphylloides* and not to other cactus-like *Euphorbia* of the Canary Islands (*E. handiensis*) and Africa. Here we aimed to reconstruct the evolutionary history of *E. canariensis* at two levels: (i) a phylogenetic approach aimed at unravelling relationships of this species and large-scale biogeographic patterns, and (ii) a phylogeographic approach focused on the history of colonization between islands of the Canarian archipelago in relation to habitat availability for this species through time. Based on previous phylogenetic studies of *Euphorbia*, we sequenced the ITS region for *E. canariensis* and several potentially related species to build a phylogenetic framework. We also sequenced two cpDNA regions for 92 individuals from 29 populations of *E. canariensis* representing its distribution range. We estimated the number of inter-island colonization events using PAICE, a recently developed method that includes a sampling effort correction. Additionally, we used species distribution modelling (SDM) to project current habitat availability for *E. canariensis* to past periods. Phylogenetic results supported the Canarian *E. canariensis* as closely related to the Southeast Asian *E. epiphylloides* and *E. sessiliflora*. In the Canarian archipelago, *E. canariensis* displayed a surprising west-to-east colonization pattern. The estimated number of inter-island colonization events was c. 20 – 50, and SDM suggested an increase in habitat availability in recent times. In summary, in this study we confirmed an extreme biogeographic disjunction between Macaronesia and Southeast Asia, described only for a small number of plant species, and estimated relatively low dispersal capabilities for *E. canariensis*, with a west-to-east colonization pattern in the Canary Islands.

**Highlights:**

- The Canarian *Euphorbia canariensis* is sister to Southeast Asian species.
- This lineage shows one of the widest disjunctions affecting Macaronesian plants.
- The lack of intermediate relatives could be the result of extinction events.
- *E. canariensis* displays low colonization ability in the Canarian archipelago.

## 1. Introduction

Oceanic islands have attracted scientific interest for centuries because they provide a clear spatiotemporal framework for evolutionary research (Darwin, 1859; Wallace, 1880; Whittaker and Fernández-Palacios, 2007). This island type is characterized by emergence from the oceanic floor without any previous contact with other land masses. Therefore, occurrence of terrestrial organisms on them can only be explained by long-distance dispersal (LDD) events (Vargas, 2014; Whittaker and Fernández-Palacios, 2007). In the case of plant species, morphological structures of diaspores have been traditionally associated with LDD processes (van der Pijl, 1982), and these LDD syndromes are frequently assumed to be responsible for oceanic island colonization (Carlquist, 1966). However, more recent studies have shown a large proportion of species unspecialized for LDD in oceanic archipelagos (Arjona et al., 2018; Heleno and Vargas, 2015; Vargas et al., 2014). In the case of the Canary Islands, one of the best-known oceanic archipelagos (Emerson and Kolm, 2005), statistical differences in island colonization between species with LDD syndromes and unspecialized species have been found (Arjona et al., 2018).

The Canary Islands are an archipelago formed by seven main islands that are the result of a hot spot in the Atlantic Ocean active since c. 60 Ma, although the oldest extant island dates from c. 20 Ma (Troll and Carracedo, 2016). Due to the closeness of the Canary Islands to the Mediterranean region and Africa throughout the geological history of the archipelago, the Canarian flora is mainly derived from those mainland regions (del Arco Aguilar and Rodríguez Delgado, 2018; Fernández-Palacios et al., 2011). One of the most remarkable species of the Canarian flora is *Euphorbia canariensis*, a major component of lowland xerophytic communities known, in Spanish, as ‘cardonal-tabaibal’ (Bramwell and Bramwell, 1974). Commonly known as ‘cardón’ in Spanish, *E. canariensis* is endemic and widely distributed in the Canarian archipelago (Arechavaleta et al., 2010). Despite the LDD process is considered critical to allow oceanic island colonization (Nathan, 2006; Traveset et al., 2014), *E. canariensis* does not display LDD syndromes. Although general patterns of island colonization can be derived from floristic studies and distribution ranges (Arjona et al., 2018; Heleno and Vargas, 2015; Vargas et al., 2012), more detailed inference of colonization patterns for a particular species can be derived from phylogeographic analyses (Coello et al., 2022, 2021). Therefore, a detailed study of inter-island colonization events is required to understand the phylogeographic history of *E. canariensis*. Furthermore, as colonization is a two-stage process that consists of dispersal and establishment (Heleno and Vargas, 2015; van der Pijl, 1982), information about habitat availability in the archipelago could help understand the colonization patterns of a species unspecialized for LDD such as *E. canariensis* (Coello et al., 2021).

*E. canariensis* is one of the two cactus-like species of *Euphorbia* in the Canary Islands together with *E. handiensis*. They are both included in *Euphorbia* subg. *Euphorbia* sect. *Euphorbia* following Bruyns et al. (2011), but *E. canariensis* appears to belong to an early diverging lineage within this group, while *E. handiensis* is part of the core of the group. It has been commonly assumed that *E. canariensis* (together with several other Canarian species) displays a Rand Flora pattern (del Arco Aguilar and Rodríguez Delgado, 2018). This is a biogeographic pattern in which multiple unrelated lineages display a similar disjunct distribution along the edges of Africa (Pokorny et al., 2015; Sanmartín et al., 2010), which can also extend to Southwest Asia and the Mediterranean Basin (e.g. *Plocama*; Rincón-Barrado et al., 2021). In particular, *Euphorbia* displays disjunctions that reach Southwest Asia, such as that of the Canarian *E. balsamifera* and its close relatives (Riina et al., 2021). The case of *E. canariensis* appears to be more extreme, as Bruyns et al. (2011) inferred that it is sister to the Southeast Asian *E. epiphylloides*, an endangered species with no more than 250 individuals in Saddle Peak, North Andaman Island (Indian Ocean) (IUCN, 1998), more than 10,000 km away from the Canarian archipelago.

Unfortunately, the relationship between the two species was not well supported (Bruyns et al., 2011), and further studies investigating phylogenetic relationships among *Euphorbia* species did not include *E. canariensis* (Dorsey et al., 2013; Horn et al., 2014). If the close relationship between *E. canariensis* and Southeast Asian species were corroborated, it would be very remarkable biogeographically given the great geographic distance between both species, although it would not be an isolated example. For example, the Canarian *Pinus canariensis* is sister to the Himalayan *P. roxburghii* (Jin et al., 2021), and Macaronesian species of *Dracaena* are closely related to species of Southeast Asia (Celiński et al., 2020; Durán et al., 2020; Edwards et al., 2018). Additionally, there are other species morphologically related to *E. epiphylloides* (i.e. the Southeast Asian *E. sessiliflora*, a species that has not yet been studied phylogenetically) that could be relevant to resolve the phylogenetic relationships of *E. canariensis*. Furthermore, if the relationship between *E. canariensis* and Asian species were confirmed, knowing the colonization abilities of this species would be useful to interpret this extreme disjunction as the result of either LDD or extinction processes in the intervening regions. To that end, it would also be relevant to estimate divergence times among species and compare them to the onset of biogeographic barriers between both territories (e.g. the aridification of Saharan, Arabian and Syrian desert; Oberprieler, 2005; Sanmartín, 2003).

In this study, we address the evolutionary history of *E. canariensis* at two levels: (i) a phylogenetic approach to clarify the relationships of *E. canariensis* and to evaluate the potential biogeographic disjunction between the Canary Islands and Southeast Asia; and (ii) a phylogeographic approach to study the pattern of colonization between islands of the Canarian archipelago in relation to habitat availability for this species through time. We hypothesized that the lack of LDD syndromes has resulted in a limited number of inter-island colonization events despite a wide potential distribution (i.e. this species occurs on every island of the archipelago except for Lanzarote). In particular, we addressed the following specific objectives: (i) to determine the phylogenetic relationships of *E. canariensis*, (ii) to calculate the divergence times among *E. canariensis* lineages and those of the sister group, (iii) to describe the geographical distribution of plastid DNA diversity of this species across the Canarian archipelago, (iv) to estimate the number of inter-island colonization events experienced by *E. canariensis* in the Canary Islands, and (v) to explore the extent of suitable habitats of this species for the present and past periods.

## 2. Material and methods

### 2.1. Study species

*Euphorbia canariensis* L. (Euphorbiaceae) is a succulent shrub, usually with 4- angled branches and curved paired spines along the branches. It looks like an up to 3-4 m tall cactus filled with latex (Bramwell and Bramwell, 2001) (Figure 1). *E. canariensis* is endemic to the Canary Islands, a volcanic archipelago located between 27.5° – 29.5° N and 13° – 18.5° W, less than 100 km off the coast of Africa. The archipelago currently comprises seven main islands that emerged from a mantle plume, in which easternmost islands are older. Current emerged islands date from >20 Ma (Fuerteventura) to c. 1 Ma (El Hierro) (Troll and Carracedo, 2016). *E. canariensis* naturally occurs on every major island of the archipelago except for Lanzarote (Arechavaleta et al., 2010). Although there are isolated historical references to its presence in Lanzarote, wild populations have not been seen on this island in recent times (Bramwell and Bramwell, 2001). The species inhabits rocky slopes, cliffs and lava fields from sea level to 900 m (Bramwell and Bramwell, 1974), and is a characteristic species of lowland xerophytic communities known, in Spanish, as ‘cardonal-tabaibal’. Regarding LDD syndromes, *E. canariensis* does not show any structure related to LDD neither in fruits nor seeds (Arjona et al., 2018).

**Figure 1:**
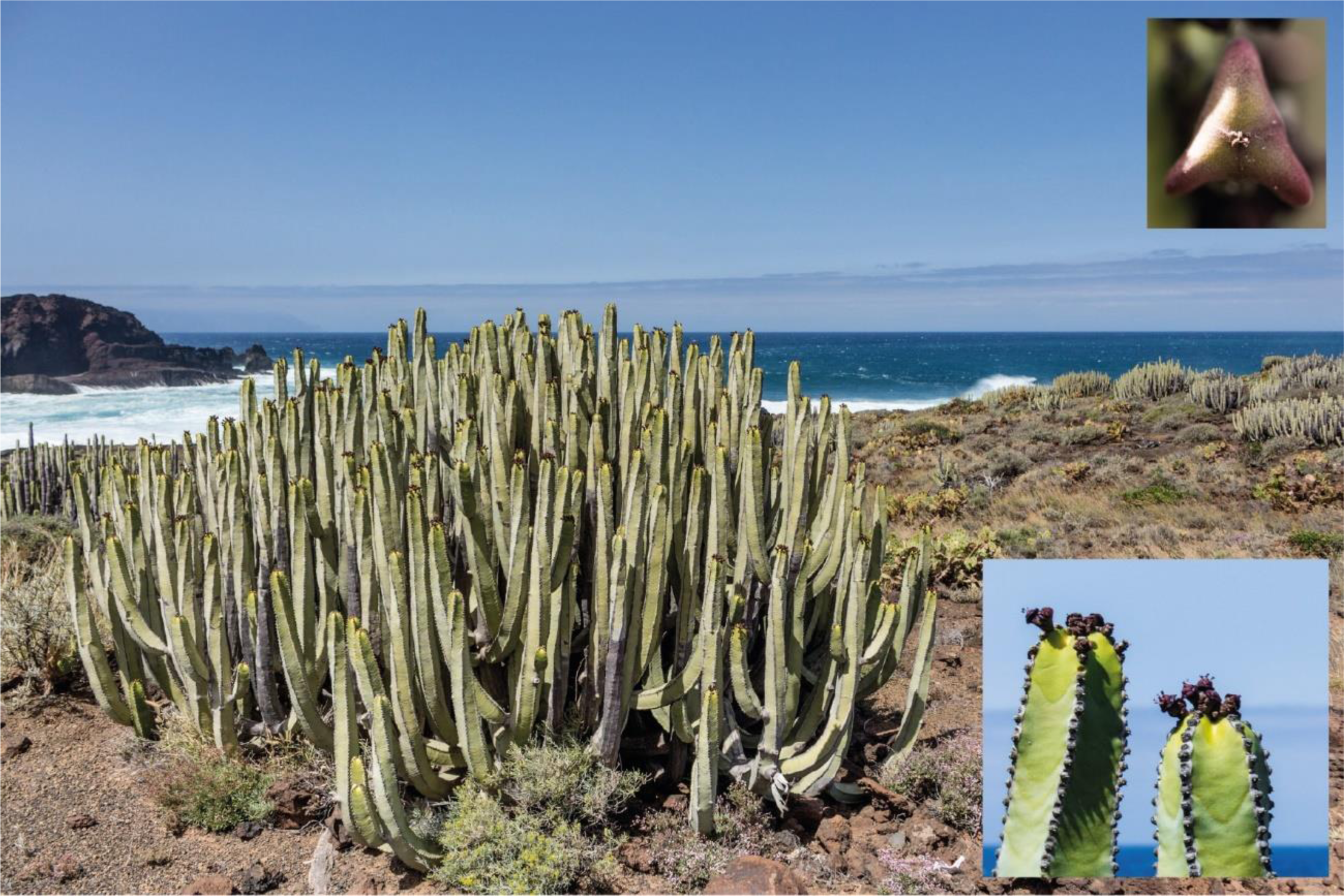
*Euphorbia canariensis* in Teno (Tenerife). Insets show fruit (top right) and flower (bottom right) details. Photos by Alberto J. Coello.

### 2.2. Sampling and DNA sequencing

We sampled fresh material of *E. canariensis* for 119 individuals from 30 populations (up to 5 individuals per population) in the six major islands of the Canarian archipelago where wild populations of the species currently occur (an old, probably cultivated individual from Lanzarote was also sampled; Table S1 in Supporting Information). Since *E. canariensis* does not develop conventional leafs, we collected an epidermal fragment from each sampled individual. We also obtained dry material from *E. abdelkuri*, *E. epiphylloides, E. handiensis, E. sessiliflora* and *E. officinarum* (Table S1 in Supporting Information). These samples were selected to get a deeper insight into the phylogenetic relationships of *E. canariensis*, in combination with sequences available in GenBank. In particular, *E. epiphylloides* has been suggested to be the most closely related species to *E. canariensis* (Bruyns et al., 2011) and they both appear to form an early diverging lineage within sect. *Euphorbia*, although this relationship needs to be corroborated. *E. abdelkuri* is also part of an early diverging lineage in sect. *Euphorbia* (Bruyns et al., 2011; Dorsey et al., 2013; Horn et al., 2014). *E. sessiliflora* (Roxburgh and Carey, 1832) has been described as morphologically similar to *E. epiphylloides*, and therefore could also be related to *E. canariensis*, but it has not been included in any phylogenetic study to date. In the case of the other cactus-like species of sect. *Euphorbia* in the Canary Islands, *E. handiendis*, we included additional samples to corroborate the lack of a close phylogenetic relationship with *E. canariensis*. Finally, *E. officinarum* was included in previous phylogenies (= *E. echinus*) and recovered as closely related to *E. handiendis*. Given the cactus-like morphology of *E. officinarum* and the geographical proximity to the Canarian archipelago (it occurs in northwestern Africa), we considered that additional material was required to corroborate its phylogenetic position.

We extracted total genomic DNA using the CTAB method (Cullings, 1992; Doyle and Doyle, 1987). For the phylogenetic analysis (see below) we sequenced the ITS region from 13 individuals of *E. canariensis* representing plastid haplotypes detected in the phylogeographic analysis (see Results) and from the other *Euphorbia* species mentioned above. For phylogeographic analysis (see below), we first selected variable cpDNA regions by performing a pilot study using 17 regions that included the most variable regions of the plastid genome of angiosperms (Shaw et al., 2007). As a result, we selected two cpDNA regions: *trn*S-*trn*G and *trn*Q-*rps*16 (Table 1). These regions were also sequenced for *E. abdelkuri*, *E. epiphylloides* and *E. handiensis* as the outgroup. Every cpDNA region was amplified by a conventional PCR in an Eppendorf Mastercycler Epgradient S. After 2 min of pretreatment at 94° C, PCR consisted of 30 – 35 cycles of 1 min at 94° C, 1 min at 52° C (48° C in some cases) and 1 min at 72° C, followed by a final elongation period of 10 min at 72° C. In every reaction, 1 µl of bovine serum albumin (BSA) at 1 mg · ml^-1^ was added to every 25 µl of reaction to improve the amplification efficiency. For problematic samples, small modifications were implemented, and internal primers were used (Table 1). PCR products were sequenced by Macrogen Inc. (Madrid, Spain), and sequences were assembled in Geneious v11.0.4 (Kearse et al., 2012).

**Table 1:**
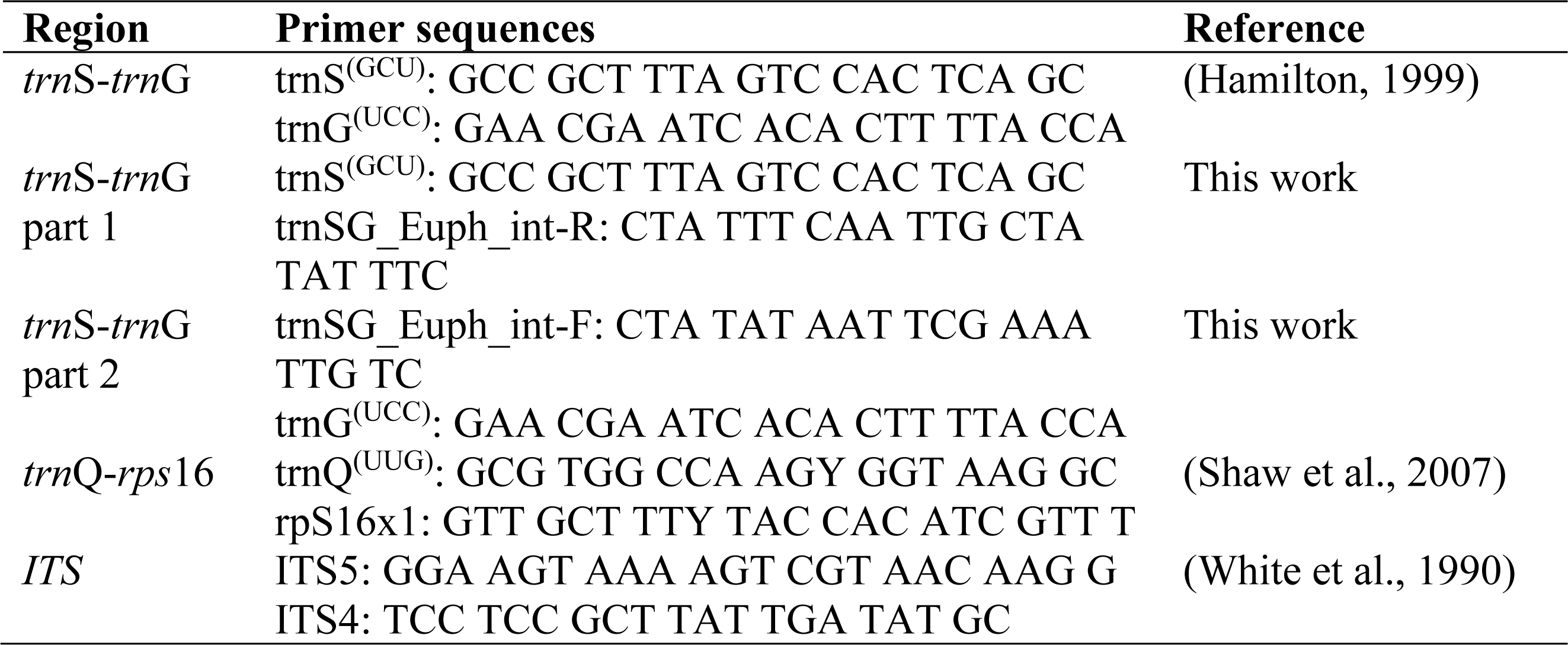
Primers used for amplification and sequencing of two cpDNA regions (trnS-trnG and trnQ-rps16) and the nDNA (ITS) in Euphorbia species.

### 2.3. Phylogenetic analysis

To analyse the phylogenetic position of *E. canariensis* and estimate divergence times, we included our new ITS sequences in the previously published alignment of 182 ITS sequences from Horn et al. (2014) (Table S2) using MAFFT v7.388 (Katoh et al., 2002). Horn et al. (2014) used 10 concatenated regions to build the most recent time-calibrated phylogeny of the genus *Euphorbia*, but we focused on the ITS region because analyses based on this nuclear DNA region alone reflect major phylogenetic patterns in the genus (and particularly in sect. *Euphorbia*) obtained by Horn *et al*.(2014). We performed a time-calibrated Bayesian phylogenetic analysis in BEAST v1.10.4 (Drummond and Rambaut, 2007) using the calibration points of Horn et al. (2014), placed at the crown nodes of the corresponding clades (Figure S1 in Supporting Information): (i) a fossil of *Hippomanoidea* used to implement an exponential prior distribution with a minimum age of 43 Ma and a mean of 2.5, (ii) a secondary calibration of the root using a normal distribution with mean 69.08 Ma and standard deviation 6.963, and (iii) a secondary calibration of Euphorbiaceae excluding Stomatocalyceae and using a normal distribution of mean 52.79 and standard deviation 5.625. We applied a GTR + I + G model of nucleotide substitution selected by the AIC criterion in jModelTest v2.1.10 (Darriba et al., 2012). An uncorrelated relaxed clock with a lognormal distribution was implemented, with a birth-death speciation process as tree prior. We executed two runs of 100 million generations each, sampled every 10,000 generations, and applied a 10% burn-in. The adequacy of both runs was analysed in Tracer v1.7.1, they were combined using LogCombiner (discarding the burn-in) and trees were summarized in a maximum clade credibility tree using median heights with TreeAnnotator.

### 2.4. Phylogeographic analysis

Sequences of the *trn*S-*trn*G and *trn*Q-*rps*16 regions from *E. canariensis* and the outgroup species *E. abdelkuri, E. epiphylloides* and *E. handiensis* (Table S1 in Supporting Information) (Bruyns et al., 2011) were aligned with MAFFT v7.388 (Katoh et al., 2002). Minor errors in the resulting alignment were corrected by visual inspection. Furthermore, a 106-bp stretch of the *trn*S-*trn*G region (from position 326 to 433 in the final alignment) was excluded for downstream analyses because preliminary exploration indicated that this fragment contained several highly homoplastic gaps and variable positions of uncertain alignment. The concatenated alignment of both regions was used to determine the genealogical relationships among haplotypes using the statistical parsimony algorithm (Templeton et al., 1992) implemented in TCS v1.21 (Clement et al., 2000). Haplotype relationships were calculated using a 95% confidence limit and gaps were treated as missing data. Additionally, to facilitate the connection between the outgroup species and *E. canariensis* haplotypes, we repeated the TCS analysis by reducing the confidence limit to 90%.

### 2.5. Estimation of inter-island colonization events

We estimated the number of inter-island colonization events based on phylogeographic data (see above) using the recently developed method PAICE (Phylogeographic Analysis of Island Colonization Events; Coello et al., 2022). We considered the palaeoisland of Mahan as a single island unit combining Lanzarote and Fuerteventura to avoid confounding within-island range expansion with inter-island colonization events, given that these two islands were connected in the Last Glacial Maximum (Troll and Carracedo, 2016) (see Results). In any case, we only included individuals from Fuerteventura, as the species does not seem to be naturally present on Lanzarote. First, we inferred the minimum number of colonization events required to explain the geographic distribution of *E. canariensis* haplotypes using the *colonization* function of the *PAICE* package (Coello et al., 2022). We also calculated asymptotic estimators of colonization events. Therefore, we built rarefaction curves for both sampling variables (genetic and field sampling), using the *rarecol* function of PAICE. We replicated rarefaction curves five times the number of levels of each variable, i.e. 140 replicates for field sampling (28 levels, one for each of the 28 populations sampled, as population was used as field sampling unit) and 75 replicates for genetic sampling (15 levels, one for each of the 14 variable positions in addition to the case of 0 variable position, i.e. not genetic information considered).

These rarefaction curves were later used to calculate the asymptotic estimators of colonization events using the *maxCol* function of *PAICE*. We deleted extreme values (i.e. those below the 2.5% quantile and above the 97.5% quantile) using argument *del = 0.05*, and calculated the 95% confidence interval of asymptotic estimators using the argument *level = 0.95*. Finally, we used the argument *method = 1* to allow the algorithm to fit the accumulation curve of colonization events with fewer parameters if curves could not be fitted initially. As the outgroup was connected to *E. canariensis* at two points (two missing haplotypes between haplotypes A and B; see Results), we repeated the analysis considering each of these possible ancestral haplotypes (see Coello et al., 2022 for additional information).

We also estimated the colonization history of *E. canariensis*, including the number of inter-island colonization events, using a Bayesian discrete phylogeographic analysis (DPA; Lemey et al., 2009) in BEAST v1.10.4 (Drummond and Rambaut, 2007) for comparison with PAICE results. We used *E. abdelkuri*, *E. epiphylloides* and *E. handiensis* as the outgroup. The *trn*S-*trn*G and *trn*Q-*rps*16 regions were used with the GTR substitution model following the AIC criterion implemented in jModelTest 2.1.10. We constrained *E. canariensis* as a monophyletic group (see Results) and implemented a uniform prior distribution for the crown age between 0.3 and 2.72 Ma, and another calibration point in a monophyletic group including *E. canariensis* and *E. epiphylloides* with a uniform prior distribution for the crown age between 3.95 and 12.68 Ma; all calibration points were obtained from the time-calibrated phylogenetic analysis described above (see Results). We used an asymmetric substitution model to reconstruct ancestral areas and a Bayesian stochastic search variable selection (BSSVS) to infer statistically supported migration routes. Six areas were defined: Mahan (palaeoisland of Fuerteventura and Lanzarote), Gran Canaria, Tenerife, La Gomera, El Hierro and La Palma. We coded continental outgroup species *E. abdelkuri*, *E. epiphylloides* and *E. handiensis* as missing data to exclude colonization of the Canarian archipelago from the continent and focus on inferring inter-island colonization events of *E. canariensis*. We used an uncorrelated relaxed clock with a lognormal distribution for the DNA partition and a strict clock for the area partition, with a simple constant size coalescent tree prior to facilitate converge. We performed three runs of 100 million generations each with a 10% burn-in, and convergence between chains was confirmed with Tracer v1.7.1. All runs were combined using LogCombiner (without burn-in) and trees were summarized in a maximum clade credibility tree using median heights in TreeAnnotator. To calculate Bayes factors (BF) of migration routes we used SpreaD3 0.9.6 (Bielejec et al., 2016), and BF values above 3 were considered as evidence for connections between areas (Kass and Raftery, 1995).

### 2.6. Species distribution modelling

For species distribution modelling (SDM), occurrences of *E. canariensis* were obtained from two sources. On the one hand, locations of populations sampled for phylogeographic analysis (Table S1 in Supporting Information) were also employed in SDM. On the other hand, we downloaded occurrences from GBIF (GBIF.org, 2021) and applied a filter to exclude those with error >1 km in their coordinates (occurrences without this information were also excluded). In order to use high quality occurrences, we selected presences from the databases Anthos, MAGRAMA, iNaturalist and observation.org. In the case of the citizen science databases (i.e. iNaturalist and observation.org), we carefully checked every occurrence that met the above requirements to validate their quality. Specifically, we only selected occurrences with attached photos in order to confirm the identification of every specimen. We also checked if the environment shown in the photo corresponded to the coordinates (e.g. we discarded observations in which the sea was close in the photos but coordinates did not correspond to coastal localities, indicating that coordinates were wrong) and whether the environment in the coordinates corresponded to the habitat of the species (e.g. we discarded occurrences at the top of Mount Teide in Tenerife). To reduce spatial bias by oversampling we followed suggestions by Boria et al. (2014) and applied a buffer of 0.05° (i.e. c. 3 arc-min). Using this procedure, we obtained a final set of 53 occurrences used in SDM (Table S3 in Supporting Information).

We downloaded 19 bioclimatic variables for current conditions from CHELSA (Karger et al., 2017) at 30 arc-seconds resolution, and 10 edaphic layers from SoilGrids (https://soilgrids.org/) at 250 m resolution (Table S4 in Supporting Information). We clipped both datasets to the extent of the study area (from 19° to 13° W and from 27° to 30° N). For edaphic layers, we reduced resolution to match that of bioclimatic layers using the *resample* function of the R package *raster* with the bilinear method (i.e. argument *method = “bilinear”*). We followed recommendations of Dormann et al. (2013) and discarded variables with |r| > 0.85 and a variance inflation factor VIF > 10. VIFs were calculated using the R package *HH* (Heiberger, 2017). This procedure resulted in a selection of eight variables: temperature seasonality (bio 4), minimum temperature of the coldest month (bio 6), precipitation seasonality (bio 15), precipitation of the driest quarter (bio 17), volumetric percentage of coarse fragments with size > 2 mm (eda 2), weight percentage of clay particles with size < 0.0002 mm (eda 3), cation exchange capacity of soil (eda 6) and pH index measured in water solution (eda 9). Climatic variables were also downloaded from PaleoClim (www.paleoclim.org) for the following past periods: Meghalayan (late-Holocene: 0.3 – 4.2 ka), Northgrippian (mid-Holocene: 4.2 – 8.326 ka), Greenlandian (early-Holocene: 8.326 – 11.7 ka), Younger Dryas Stadial (11.7 – 12.9 ka), Bølling-Allerød (12.9 – 14.7 ka), Heinrich Stadial 1 (14.7 – 17.0 ka), Last Glacial Maximum (c. 22 ka) and Last Interglacial (c. 130 ka). Three Pleistocene climatic periods (MIS19, ca. 787 ka; mid-Pliocene warm period, 3.205 Ma; and M2, ca. 3.3 Ma), were not used because not all bioclimatic variables were available for them. We assumed that edaphic variables were constant throughout all periods and therefore used the same edaphic layers for every period (they were resampled to match the extension and resolution of climatic layers).

We estimated the current distribution model of *E. canariensis* by employing the maximum entropy algorithm implemented in Maxent 3.4.1 (Phillips et al., 2006), as it performs well when using presence-only data (Elith et al., 2006). We used 60% of occurrences for training the algorithm and the other 40% of occurrences for testing. We conducted 100 sub-sample replicates. The model based on current conditions was projected to past periods to reconstruct the potential distribution of this species through time.

## 3. Results

### 3.1. Phylogenetic relationships of *E. canariensis*

The ITS phylogeny (Figure 2 and Figure S1 in Supporting Information; Alignment S1 in Supporting Information) confirmed that both cactus-like species of *Euphorbia* occurring in the Canary Islands (*E. canariensis* and *E. handiensis*) are not sister species (Figure 2). In the case of *E. canariensis*, sequenced individuals (at least one per islands and from different haplotypes, see above) formed a monophyletic group (PP = 1, 0.3 – 2.72 Ma), which was grouped with the Southeast Asian *E. epiphylloides* and *E. sessiliflora* (PP = 0.97, 3.95 – 12.68 Ma). This clade was suggested to be closely related to *E. abdelkuri* but with low support (PP = 0.55) and had an early-diverging position within sect. *Euphorbia*. In contrast, *E. handiensis* (PP = 0.73, 0 – 0.92 Ma) was located in the core of sect. *Euphorbia*, sister to *E. officinarum* (PP = 1, 0.16 – 2 Ma) from Northwest Africa.

**Figure 2:**
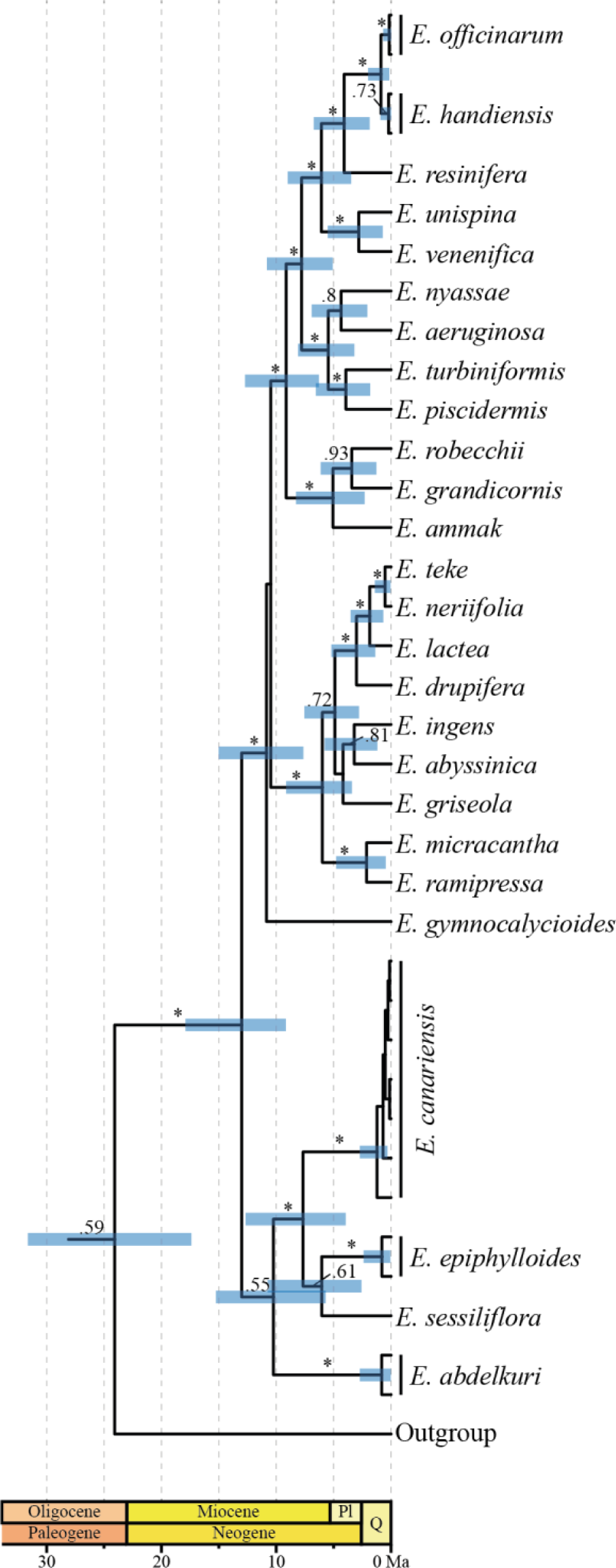
Time-calibrated Bayesian phylogenetic analysis of *Euphorbia* subg. *Euphorbia* sect. *Euphorbia* (the complete phylogeny is displayed in Figure S1 in Supporting Information). Numbers above branches indicate posterior probabilities (PP) when PP > 0.5 and asterisks (*) indicate PP > 0.95. Bars at nodes which PP > 0.5 indicate the 95% highest posterior density intervals for ages.

### 3.2. Phylogeographic analysis

We obtained DNA sequences of the *trn*S-*trn*G and *trn*Q-*rps*16 plastid DNA regions for 92 individuals from 29 populations of *E. canariensis* (Table S1 in Supporting Information). However, the individual from Los Canarios (La Palma) could not be sequenced for one of the two regions and this population was only used for SDM. Concatenation of *trn*S-*trn*G and *trn*Q-*rps*16 sequences of *E. canariensis* and the outgroup resulted in an alignment of 1108 bp (565 bp for *trn*S-*trn*G and 543 bp for *trn*Q-*rps*16; Alignment S2 in Supporting Information). We observed 13 variable positions, and the TCS analysis resulted in 10 substitution-based haplotypes for *E. canariensis* (Figure 3). The most frequent haplotype was I, observed in 47 of 92 individuals (51%). The haplotype network showed no loops, few (four) missing haplotypes, and three main groups of haplotypes: one group endemic to La Gomera (haplotype A); a second group found on the westernmost islands (La Palma and El Hierro; haplotypes B and C); and a third group (containing seven haplotypes, D-J) occurring on central and eastern islands (Tenerife, Gran Canaria and Fuerteventura). The probably cultivated individual from Lanzarote displayed haplotype I, the most common haplotype, also found in Fuerteventura (the other part of the Mahan palaeoisland). Tenerife had the highest number of haplotypes (five) while La Gomera had the lowest (only one haplotype). The outgroup species (*E. abdelkuri*, *E. epiphylloides* and *E. handiensis*) were not connected to *E. canariensis* haplotypes at a 95% confidence interval, but they were connected to the two missing haplotypes between A and B at a 90% confidence interval.

**Figure 3:**
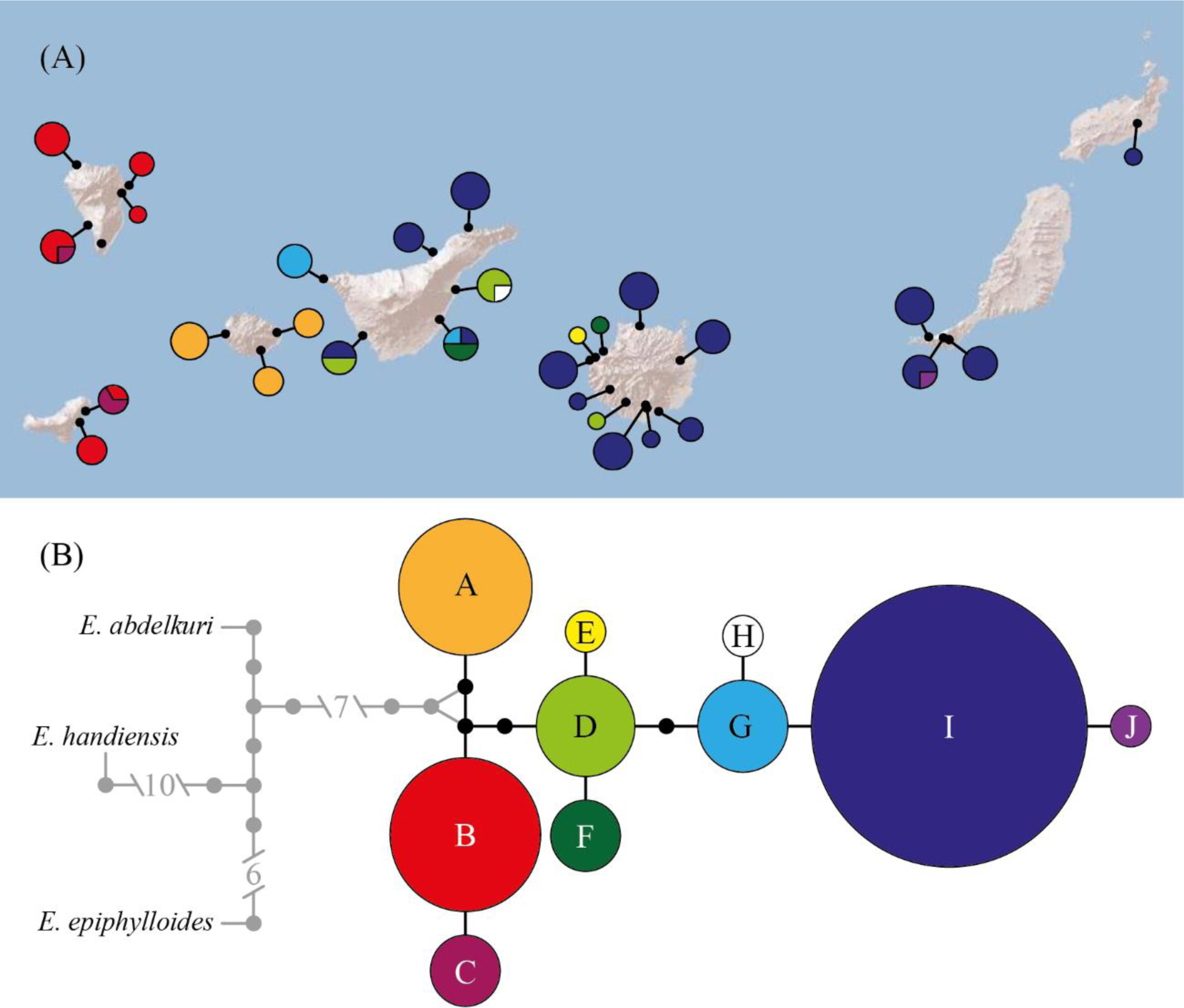
Phylogeographic analysis of *Euphorbia canariensis*. **A:** Geographic distribution of haplotypes of *E. canariensis* based on two cpDNA regions (*trn*S-*trn*G and *trn*Q-*rps*16). Circle sizes are proportional to the number of individuals (note that the southernmost population of La Palma could not be sequenced). **B:** statistical parsimony network of *E. canariensis* haplotypes; lines correspond to single nucleotide substitutions, dots indicate missing haplotypes (extinct or not found), and circle sizes are proportional to haplotype frequencies; the outgroup, in grey, is only connected to *E. canariensis* haplotypes at lower confidence intervals (see text).

### 3.3. Estimation of inter-island colonization events

According to PAICE results, haplotypes of *E. canariensis* require a minimum of eight colonization events between islands of the Canarian archipelago to explain their geographic distribution and genealogical relationships. When sampling bias was considered, rarefaction curves fitted well to theoretical curves (Figure 4) and the asymptotic estimators for the number of colonization events were as follows: (i) when using the missing haplotype closest to haplotype A as ancestral, 22.5 (95% CI: 20.6 – 25.1) colonization events for the genetic estimator and 28.1 (95% CI: 23.9 – 34.5) colonization events for the field estimator; and (ii) when using the missing haplotype closest to haplotype B as ancestral, 26.4 (95% CI: 22.8 – 32.7) colonization events for the genetic estimator and 38.3 (95% CI: 31.4 – 49.7) colonization events for the field estimator.

**Figure 4:**
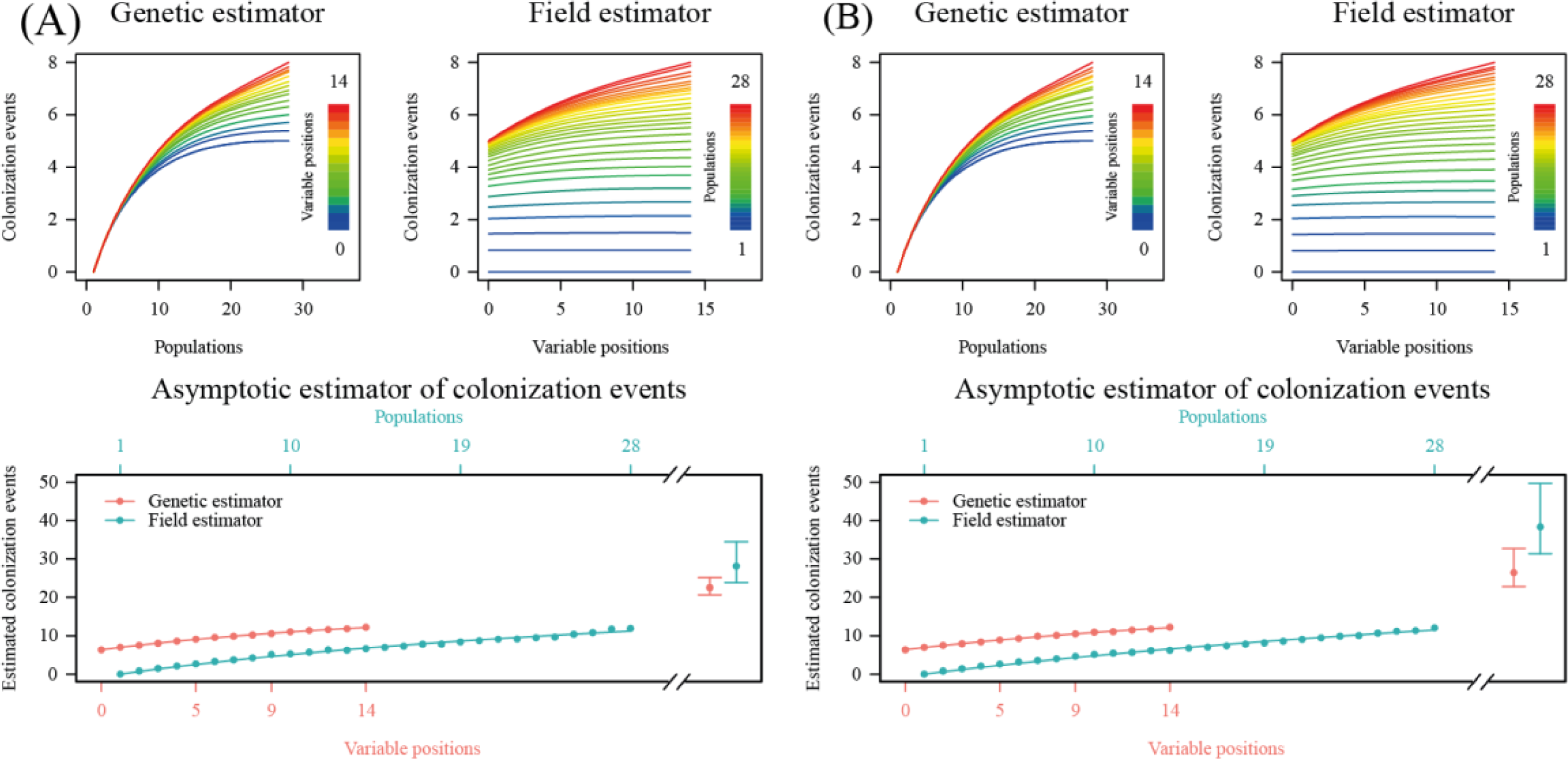
Rarefaction curves of colonization events of *E. canariensis* in the Canary Islands. **A:** Results when considering the missing haplotype closest to A as ancestral. **B:** Results when considering the missing haplotype closest to B as ancestral. Top plots represent raw rarefaction curves of both genetic (left) and field (right) estimators, and bottom plots show rarefaction curves of the final estimation for both genetic (red) and field (blue) estimators, in which asymptotic estimators are plotted with their 95% confidence intervals.

Bayesian discrete phylogeographic analysis (DPA) estimated 13.3 (95% CI: 9 – 19) colonization events. The estimated divergence time between *E. canariensis* and *E. epiphylloides* (PP = 1) was 3.95 – 9.14 Ma, while the estimated age for the most recent common ancestor of *E. canariensis* haplotypes was 1.76 – 2.72 Ma (PP = 1) (Figure 5). La Gomera was suggested as the most probable ancestral area of *E. canariensis* in the Canarian archipelago (45%), followed by Tenerife (30%). From this common ancestor, three well supported clades were differentiated, including two western clades (haplotype A, with La Gomera as the most probable ancestral area, 100%); and haplotypes B-C, with La Palma as the most probable ancestral area (68%) and subsequent colonization of El Hierro); and an eastern clade (haplotypes D-J), with Tenerife as the most probable ancestral area (90%) and subsequent colonization of Gran Canaria and Mahan. The Bayesian stochastic search variable selection (BSSVS) showed a general pattern of west to east colonization of the Canarian archipelago. The most probable routes of colonization of *E. canariensis* are shown in Table 2.

**Figure 5:**
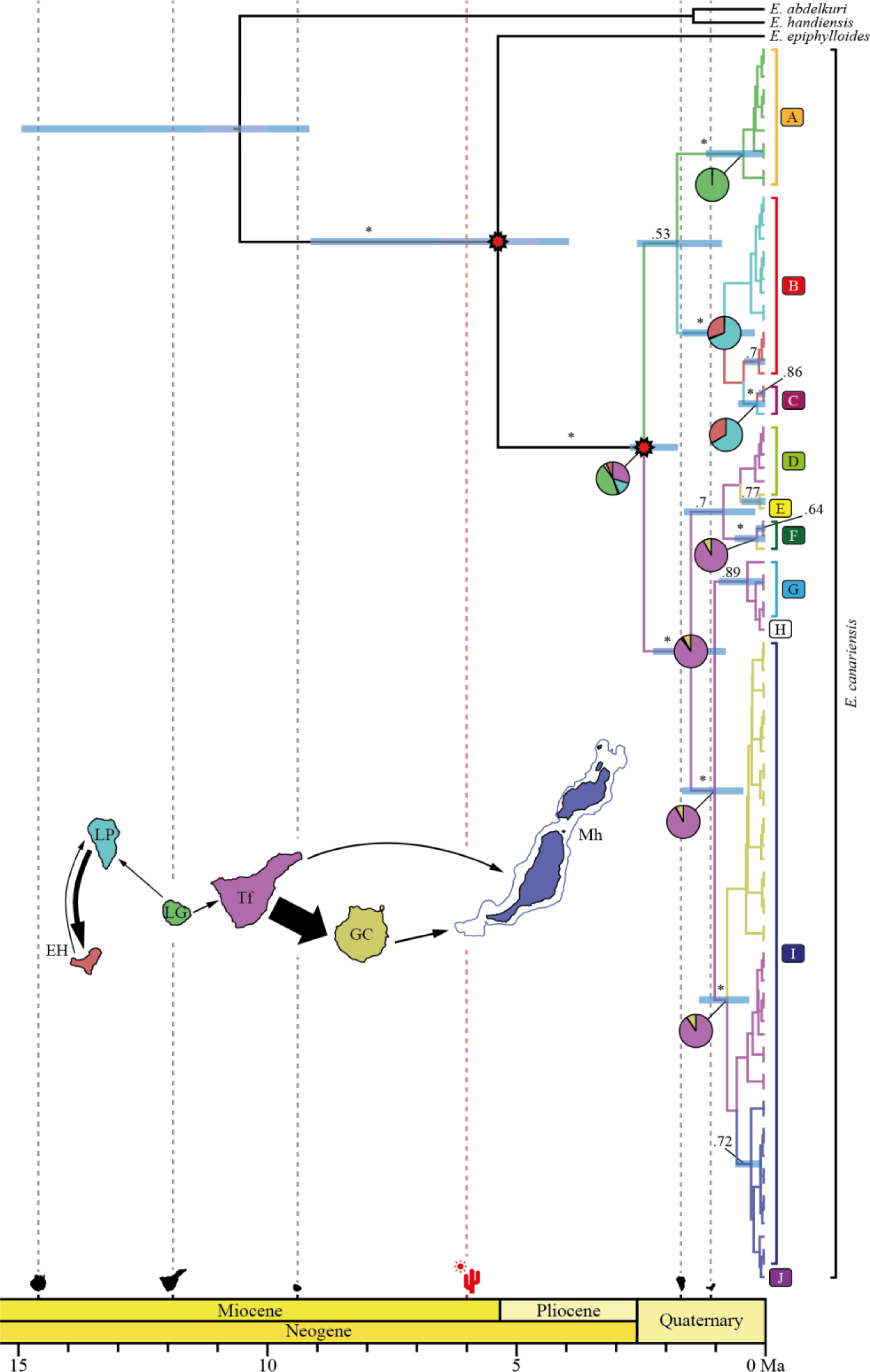
Discrete phylogeographic analysis and haplotype divergence times of *E. canariensis* based on two cpDNA regions (*trn*S-*trn*G and *trn*Q-*rps*16). Letters and colours of terminal labels in *E. canariensis* represent haplotypes as in Figure 3. Phylogenetic relationships correspond to the maximum clade credibility tree. Branch colours correspond to the most probable ancestral area as shown in the map: El Hierro (EH) in red, La Palma (LP) in light blue, La Gomera (LG) in green, Tenerife (Tf) in purple, Gran Canaria (GC) in yellow and Mahan (Mh, palaeoisland comprising Lanzarote and Fuerteventura) in dark blue (outgroup branches are not coloured). Numbers above branches are posterior probabilities (PP), only shown when PP>0.5; asterisks (*) indicate PP>0.95. Node bars indicate the 95% highest posterior density intervals for ages when PP>0.5. Pie charts indicate the posterior probability distributions of ancestral areas for well supported clades (PP>0.95) for nodes corresponding to *E. canariensis*. Red starts indicate secondary calibration points according to ages shown in Figure 2. Ages of emergence of five islands of the Canarian archipelago are indicated (GC: 14.6 Ma, Tf: 11.9 Ma, LG: 9.4 Ma, LP: 1.7 Ma, EH: 1.1 Ma; Mh is older than 20 Ma and therefore it is not shown along the time scale; Troll and Carracedo, 2016). The red dashed line at 6 Ma indicates the origin of the Saharan, Arabian and Syrian deserts (Oberprieler, 2005; Sanmartín, 2003). The inset map represents the most probable migration routes, i.e. those with Bayes factor (BF) >3; thickness of arrows is proportional to BF values.

**Table 2:** Most probable routes of colonization of *E. canariensis* in the Canary Islands according to Bayesian stochastic search variable selection (BSSVS). Only routes of colonization with a Bayes Factor (BF) greater than 3 are shown.

| **From** | **To** | **Bayes Factor (BF)** |
| --- | --- | --- |
| Tenerife | Gran Canaria | 74.82 |
| La Palma | El Hierro | 15.75 |
| Gran Canaria | Mahan | 6.19 |
| Tenerife | Mahan | 5.31 |
| La Gomera | Tenerife | 4.60 |
| El Hierro | La Palma | 3.83 |
| La Gomera | La Palma | 3.56 |

### 3.4. Species Distribution Modelling

The inferred potential distribution of *E. canariensis* for the present and past periods is shown in Figure 6. Current potential distribution of this species is located mainly in the lowlands of every island of the Canarian archipelago. The narrowest potential habitats available (relative to island size) were found on the easternmost islands (Lanzarote and Fuerteventura) while the other islands showed wider habitat availability (Gran Canaria, Tenerife, La Gomera, La Palma and El Hierro). The predictive power of the model was supported by an AUC = 0.871, and the two variables that contributed the most to the model were the minimum temperature of the coldest month (bio 6) and the pH index measured in water solution (eda 9) (Table 3). Projections displayed a tendency to narrower habitat availability in the past, with the narrowest areas in the Hainrich Stadial 1 (14.7 – 17.0 ka) and Last Glacial Maximum (c. 22 ka). The oldest period studied (Last Interglacial, c. 130 ka) did not follow this pattern, and showed a large potential distribution for this species in the Canarian archipelago.

**Figure 6:**
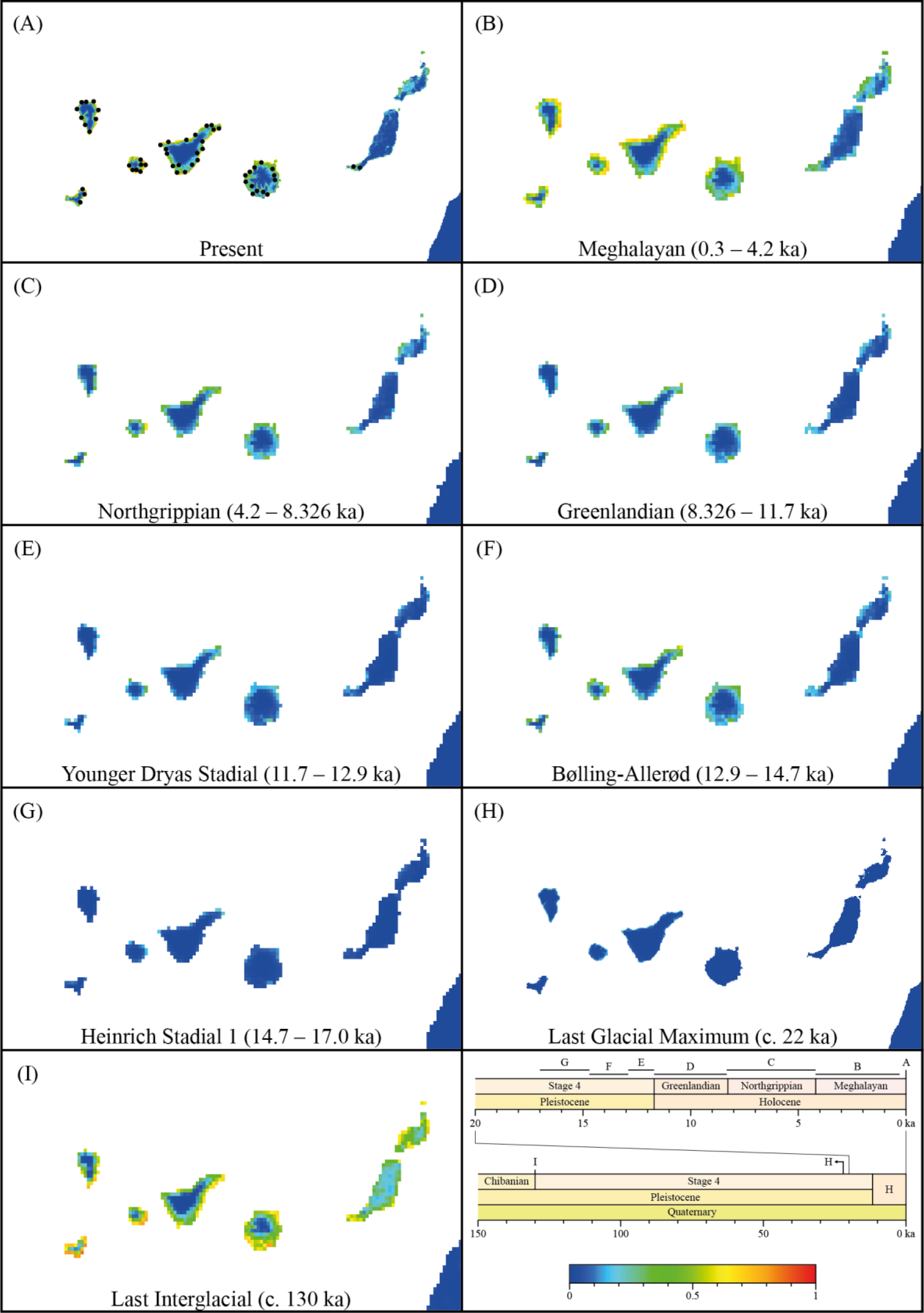
Habitat availability for *E. canariensis* in the Canary Islands through time based on species distribution modelling (SDM). **A:** Inferred suitability for the present; dots indicate occurrences used in the SDM analysis (see text); **B:** projected distribution for the Meghalayan (0.3 – 4.2 ka); **C:** projected distribution for the Northgrippian (4.2 – 8.326 ka); **D:** projected distribution for the Greenlandian (8.326 – 11.7 ka); **E:** projected distribution for the Younger Dryas Stadial (11.7 – 12.9 ka); **F:** projected distribution for the Bølling-Allerød (12.9 – 14.7 ka); **G:** projected distribution for the Heinrich Stadial 1 (14.7 – 17.0 ka); **H:** projected distribution for the Last Glacial Maximum (c. 22 ka); **I:** projected distribution for the Last Interglacial (c. 130 ka). In all maps, suitability is indicated as colours according to the scale shown at the bottom right. A time scale for the past periods used in projections is also shown.

**Table 3:** Variable contributions to the SDM analysis of E. canariensis. Variables are sorted by contribution percentage.

| **Abbreviation** | **Variable** | **Contribution (%)** |
| --- | --- | --- |
| Bio 6 | Minimum temperature of the coldest month | 46.6 |
| Eda 9 | pH index measured in water solution | 31.8 |
| Bio 17 | Precipitation of the driest quarter | 13.6 |
| Eda 2 | Volumetric percentage of coarse fragments with size > 2 mm | 3.3 |
| Bio 15 | Precipitation seasonality | 2.6 |
| Eda 3 | Weight percentage of clay particles with size < 0.0002 mm | 1.4 |
| Eda 6 | Cation exchange capacity of soil | 0.3 |
| Bio 4 | Temperature seasonality | 0.3 |

## 4. Discussion

This study supports the hypothesis of poor dispersal and colonization abilities of *E. canariensis*. The phylogenetic position of *E. canariensis* as sister to Southeast Asian species indicates a disjunction pattern wider than that of the Rand Flora, which is observed in several Macaronesian species (Pokorny et al., 2015; Sanmartín et al., 2010). The coincidence of the divergence time between *E. canariensis* and its Asian closest relatives (Figure 5) and the origin of deserts separating them (Oberprieler, 2005; Sanmartín, 2003) is consistent with extinction of intermediate lineages due to ongoing aridification in northern Africa and southwestern Asia rather than long-distance dispersal. In fact, the lack of LDD syndromes in *E. canariensis* is mirrored by a relatively low number of inter-island colonization events in comparison with other Canarian species (Coello et al., 2021).

### 4.1. An extreme disjunction between the Canary Islands and Southeast Asia

The results of the present study confirm the pattern suggested by Bruyns et al. (2011), in which *E. canariensis* is located in an early diverging lineage of *Euphorbia* subg. *Euphorbia* sect. *Euphorbia* (Figure 2). We recovered a well-supported clade including *E. canariensis* and *E. epiphylloides*, but a third species that had not yet been analyzed (*E. sessiliflora*) was added to this clade (Figure 2). In fact, morphology of *E. sessiliflora* is similar to that of *E. epiphylloides*, and both species occur in Southeast Asia, more than 10,000 km away from the Canarian archipelago. This leaves an extremely wide area between them, including the northern African continent and Southwest Asia.

It has been suggested that *E. canariensis* conforms to a Rand Flora biogeographic pattern (del Arco Aguilar and Rodríguez Delgado, 2018). This is a disjunction pattern in which sister taxa occur in peripheral regions of the African continent and adjacent islands (Pokorny et al., 2015; Sanmartín et al., 2010) and it has been described for several species of the Canarian archipelago. However, the apparent lack of any close relatives to *E. canariensis* in Africa (Figure 2; Bruyns et al., 2011) suggests that the disjunction pattern of this species goes beyond the Rand Flora. In fact, other Canarian species display a similarly extreme disjunction pattern between Macaronesia and the Southeast Asia. For example, Macaronesian dragon tree species (*Dracaena draco* from the Canary Islands, Cape Verde and Southwest Morocco; and *D. tamaranae* from Gran Canaria in the Canary Islands) diverged c. 15 Ma from their closest relatives *D. cambodiana* and *D. cochinchinensis*, both of them distributed in Southeast Asia (Celiński et al., 2020; Durán et al., 2020; Edwards et al., 2018). Similarly, in the genus *Pinus*, the Canarian pine (*P. canariensis*) is sister to the Himalayan *P. roxburghii*, and they diverged c. 15 – 25 Ma (Jin et al., 2021).

One of the hypotheses proposed to explain the Rand Flora pattern is that extant related lineages occurring in the margins of Africa are remnants of more widely distributed African lineages affected by extinction (i.e. the vicariance hypothesis; Sanmartín et al., 2010). Considering that the estimated divergence time between *E. canariensis* and its sister species (3.95 – 12.68 Ma; Figure 2) happened in a time window similar to that of the origin of the Saharan, Arabian and Syrian deserts 6 Ma (Oberprieler, 2005; Sanmartín, 2003), it is reasonable to hypothesize that extinction of intermediate lineages in the Arabian Peninsula and Africa resulted in this extreme disjunction pattern, rather than a long-distance dispersal event. For *Pinus* spp. (Kvaček et al., 2014) and *Dracaena* spp. (Denk et al., 2014), intermediate fossils support a wider past distribution followed by extinction, resulting in wide Macaronesian-Asian disjunctions. However, fossils related to the *E. canariensis* lineage have not yet been found in Africa, Asia or elsewhere. Therefore, deeper paleobotanical research might provide additional support for the extinction hypothesis.

### 4.2. *E. canariensis* in the Canary Islands

In the Canary Islands, *E. canariensis* is not the only cactus-like species of *Euphorbia*, as *E. handiensis* is endemic to the Jandía Peninsula of Fuerteventura. Phylogenetic results indicate that both species belong to different lineages of sect. *Euphorbia* (Figure 2). While *E. canariensis* is related to Southeast Asian species, *E. handiensis* is sister to the African *E. officinarum* (Figure 2; Bruyns et al., 2011). Therefore, the two species are the result of independent events of colonization of the Canarian archipelago<u>, as found in several other genera</u>.

The relatively recent divergence among *E. canariensis* haplotypes in the Canary Islands (crown age, 1.76 – 2.27 Ma; Figure 5) contrasts with the old divergence between this species and its close relatives (stem age, 3.95 – 12.68 Ma; Figure 2). This implies a high uncertainty on the time of colonization of the Canary Islands from the continent, which may have happened at any point between the stem age and the crown age (Martín-Hernanz et al., 2023). Indeed, the geographic and genetic distance between *E. canariensis* and its closest relatives highlights the problem of incomplete taxon sampling (due to extinction or unsampled close relatives). It is possible that the colonization of the Canarian archipelago from the continent is much older than the crown age if *E. canariensis* suffered significant extinction in past periods in the Canary Islands (García-Verdugo et al., 2019b), but a relatively recent colonization is also possible if significant extinction of related lineages has occurred in the continent.

In oceanic archipelagos, it is common to observe lineages of organisms whose phylogeographic relationships are concordant with the geological sequence of emergence of oceanic islands. This pattern is known as the “progression rule”, according to which a lineage of organisms colonizes islands of an oceanic archipelago as they emerge (Shaw and Gillespie, 2016). In the case of the Canary Islands, the hotspot responsible for the emergence of islands has historically followed an east-to-west trajectory, and therefore eastern islands are older (Troll and Carracedo, 2016). As a result, some species show an east-to-west pattern of lineage divergence (e.g. Villa-Machío et al., 2020), although the Canarian archipelago as a whole provides mixed evidence for and against this “progression rule” (Shaw and Gillespie, 2016). In the case of *E. canariensis*, a pattern opposite to the “progression rule” is found (Figures 3 and 5), in which central-western islands were apparently colonized earlier than eastern islands in the evolutionary history of the species when considering extant lineages. It was recently proposed by García-Verdugo et al. (2019a) that some lineages in the easternmost islands of the Canarian archipelago (Fuerteventura and Lanzarote) could be the result of recent colonization events. Therefore, the apparent lack of wild population of *E. canariensis* in Lanzarote (only one individual observed in 1990; Bramwell and Bramwell, 2001) together with its scarcity in Fuerteventura and the limited habitat availability in these eastern islands through time (Figure 6) could be related to a recent eastward colonization of these islands.

### 4.3. Colonization ability of *E. canariensis*

Our phylogeographic reconstruction and the most probable colonization routes suggest an overall west-to-east colonization pattern for *E. canariensis* (Figure 5). Given the occurrence of this species in the six major islands of the Canarian archipelago (considering Lanzarote and Fuerteventura as the single palaeoisland of Mahan; Troll and Carracedo, 2016), *E. canariensis* has experienced at least five inter-islands colonization events based on chorology alone (Arjona et al., 2018; Vargas et al., 2015). The geographic distribution of plastid DNA diversity (Figure 3) indicates a minimum of eight colonization events, and estimates obtained in PAICE considering sampling effort (Coello et al., 2022) suggested c. 20 – 50 colonization events (see Results, Figure 4). Results of this study corroborate results of Coello et al. (2022) regarding the fact that DPA analysis (c. 13 colonization events with a 95% confidence interval of 9 – 19 colonization events) estimates a smaller number of inter-island colonization events than PAICE. Additionally, the congruence between haplotype divergence times and the emergence of La Palma and El Hierro is remarkable considering that calibrations based on island ages were not implemented (see Methods). As shown in Figure 5, the clade composed of haplotypes B and C (occurring on La Palma and El Hierro) diverged from its sister haplotype (A) 0.88 – 2.58 Ma (although this relationship has low support; PP = 0.53), around the time of emergence of La Palma (1.7 Ma; Troll and Carracedo, 2016), and this island was inferred as ancestral for this clade according to the DPA. Similarly, the colonization of El Hierro by clade B – C was estimated to have happened not earlier than 1.67 Ma, coinciding with the emergence of this island (1.1 Ma; Troll and Carracedo, 2016).

In agreement with the lack of LDD syndromes in *E. canariensis* (Arjona et al., 2018), a relatively low number of inter-island colonization events is estimated for this species when accounting for sampling effort. According to the divergence times of haplotypes, *E. canariensis* underwent this number of inter-island colonization events in the Canarian archipelago in the last 1.73 – 2.72 Ma (Figure 5). Another species without LDD syndromes, *Cistus monspeliensis*, experienced c. 25 – 33 inter-island colonization events (Coello et al., 2022) in less than 1 Ma (Coello et al., 2021; Fernández-Mazuecos and Vargas, 2011) and occurs on one less island (as *C. monspeliensis* does not occur on Mahan; Arechavaleta et al., 2010). Furthermore, in Coello et al. (2022) it was estimated that one lineage of *Olea europaea* subsp. *guanchica* (displaying zoochorous drupes and occurring on one less island than *C. monspeliensis*) underwent c. 20 – 46 inter-island colonization events (Arechavaleta et al., 2010) in a shorter time than *E. canariensis* (Besnard et al., 2009). Considering that *E. canariensis* occurs on a higher number of islands than *C. monspeliensis* and *O. europaea* subsp. *guanchica*, that *E. canariensis* probably colonized the archipelago earlier (Figure 5), and that it also displays wider habitat availability (Figure 6), it seems reasonable to think that *E. canariensis* has lower colonization abilities than *C. monspeliensis* and *O. europaea* subsp. *guanchica*. In fact, this apparently low colonization capacity of *E. canariensis* remarks the improbability of a LDD event from Southeast Asia to the Canarian archipelago in the clade including *E. canariensis*, *E. epiphylloides* and *E. sessiliflora* (Figure 2), and supports extinction in intermediate locations during the evolutionary history of this clade.

## 5. Conclusions

In summary, this study supports that the Canarian *E. canariensis* is closely related to Southeast Asian species, and therefore represents one of the widest biogeographic disjunctions among Macaronesian plant species. This fact, together with the apparently low colonization capabilities of *E. canariensis* (considering the lack of LDD syndromes and the relatively low estimated number of inter-island colonization events despite a relatively old age), supports the idea of extinction of intermediate relatives of *E. canariensis* in Africa and southwestern Asia. This extreme Canarian-Asian disjunction deserves exploration in other plant groups, such as *Apollonias* and *Bosea* (Li et al., 2011; Martín-Hernanz et al., 2023).

## Declaration of Competing Interest

The authors declare that they have no known competing financial interests or personal relationships that could have appeared to influence the work reported in this paper.

## Acknowledgements

The authors thank Sara Martín-Hernanz for her continuous help about the Canarian-Asian disjunction; Brent Emerson, Alfredo García and Luis Valente for their commentaries to improve the text and the discussion; Yurena Arjona and Miguel Blázquez for sample collection; and Itziar Cantero for laboratory assistance. Collection permits were granted by the Cabildos Insulares of the Canary Islands. This study is part of the projects CGL2015-67865-P and PGC2018-101650-B-I00, funded by the Spanish Ministry of Economy, Industry and Competitiveness. A.J.C. was supported by the Spanish Ministry of Education, Culture and Sport through an FPU fellowship (reference FPU16/05681). M.F.-M. was supported by the Spanish Ministry of Economy and Competitiveness through a Juan de la Cierva fellowship (reference IJCI-2015-23459), and by the Spanish National Research Council (CSIC) through a Special Intramural Project (reference 201930E078).

## Supporting information

Additional supporting information can be found in the online version of this article:

**Table S1:**
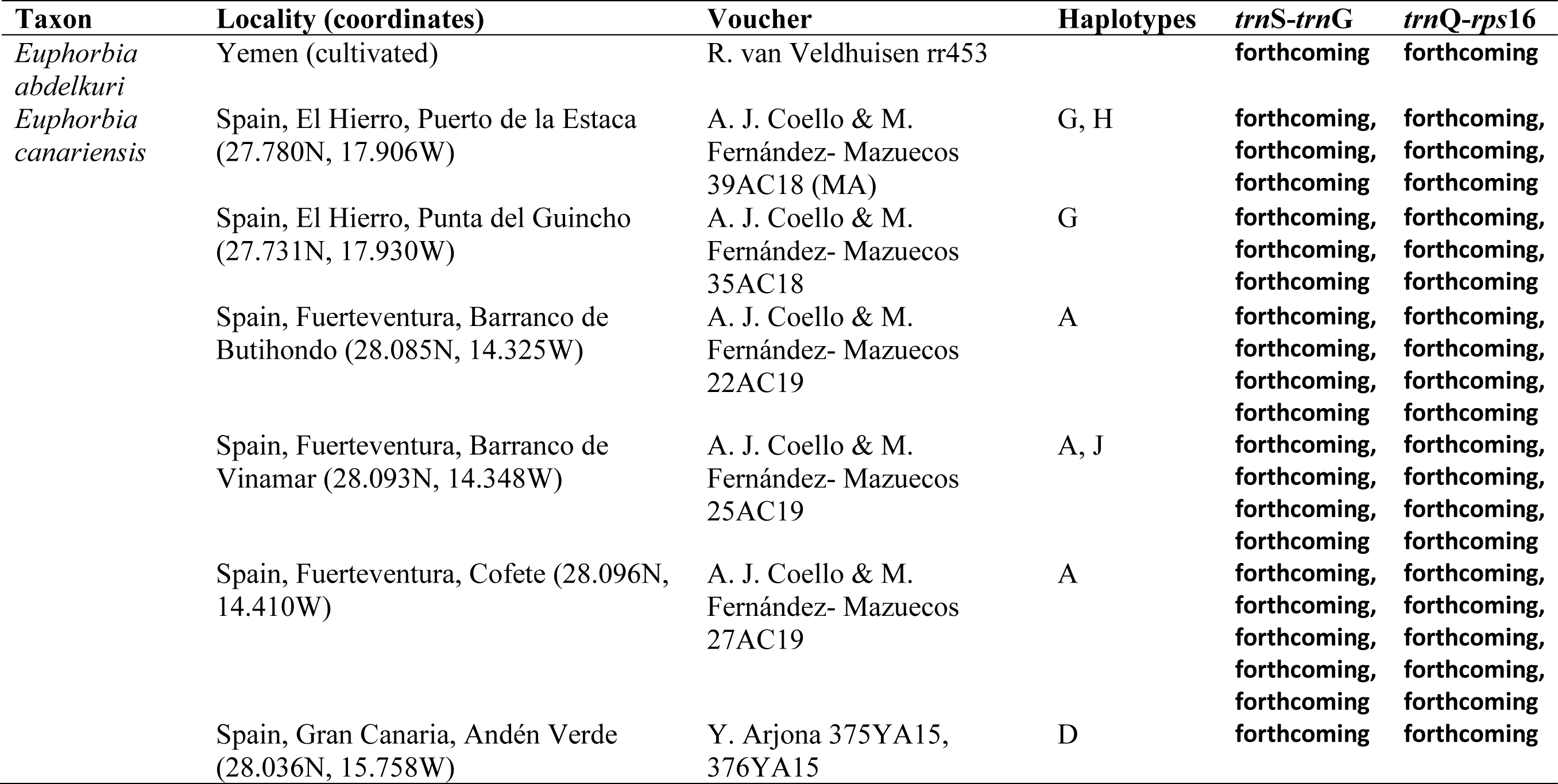
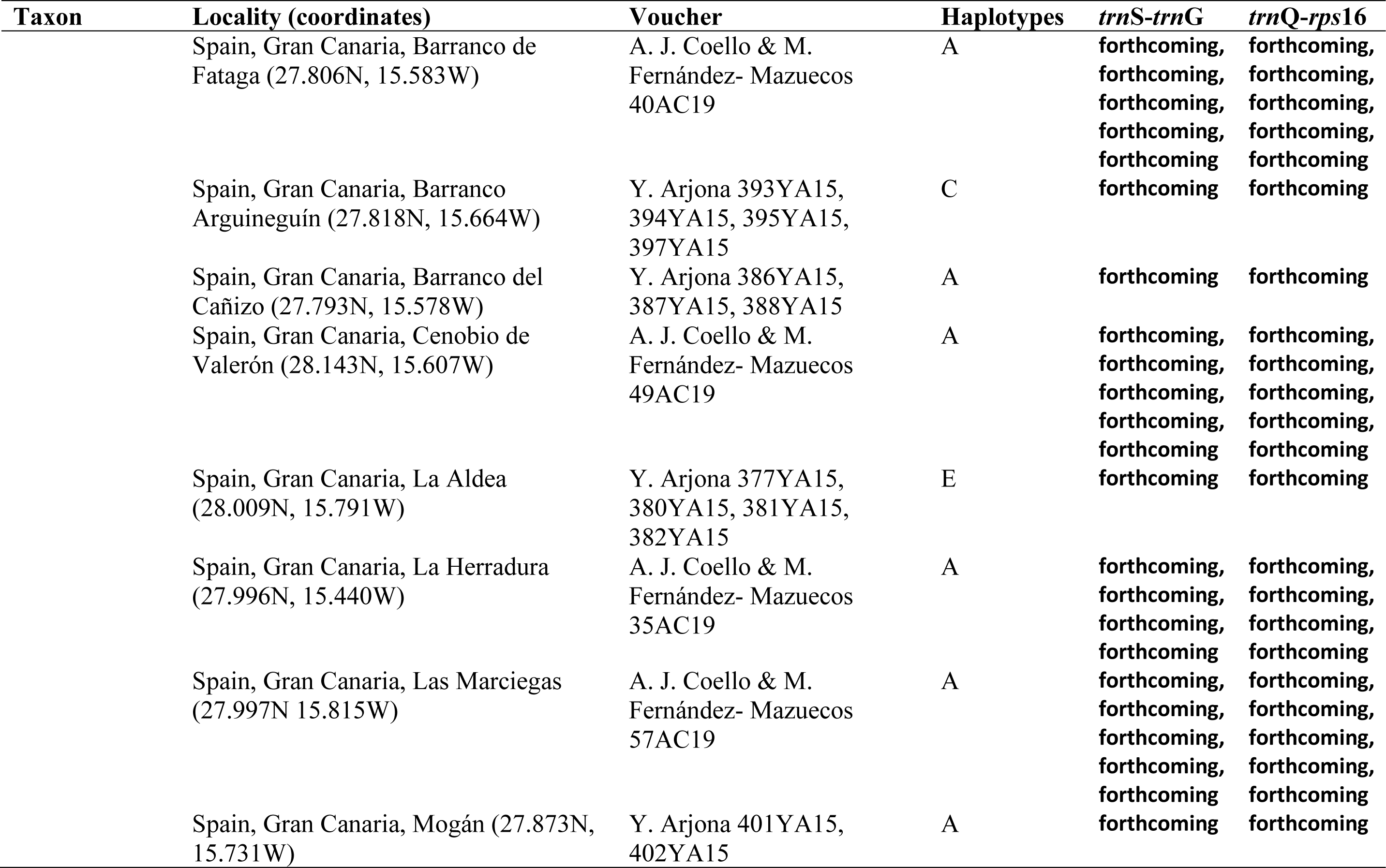
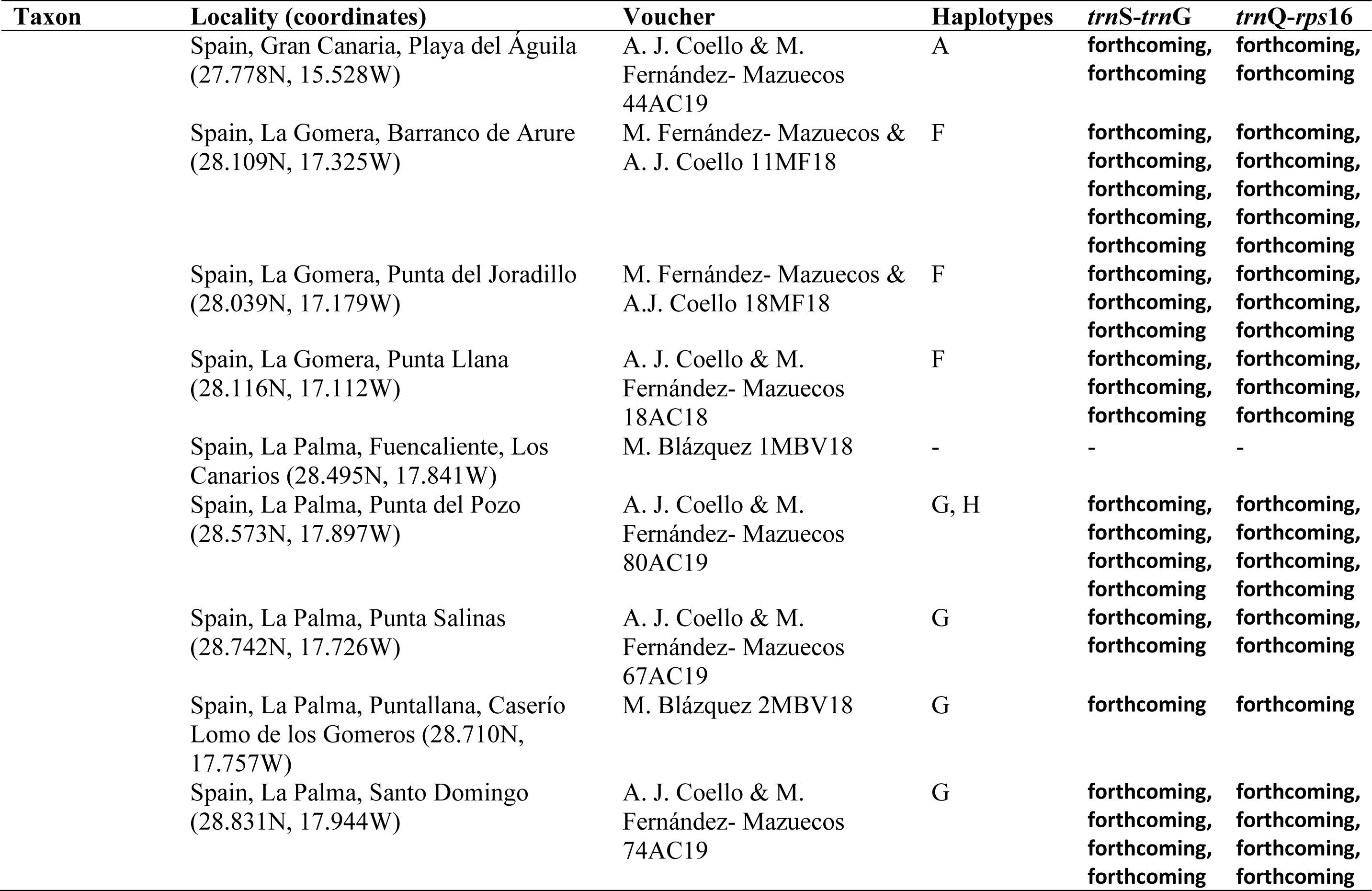
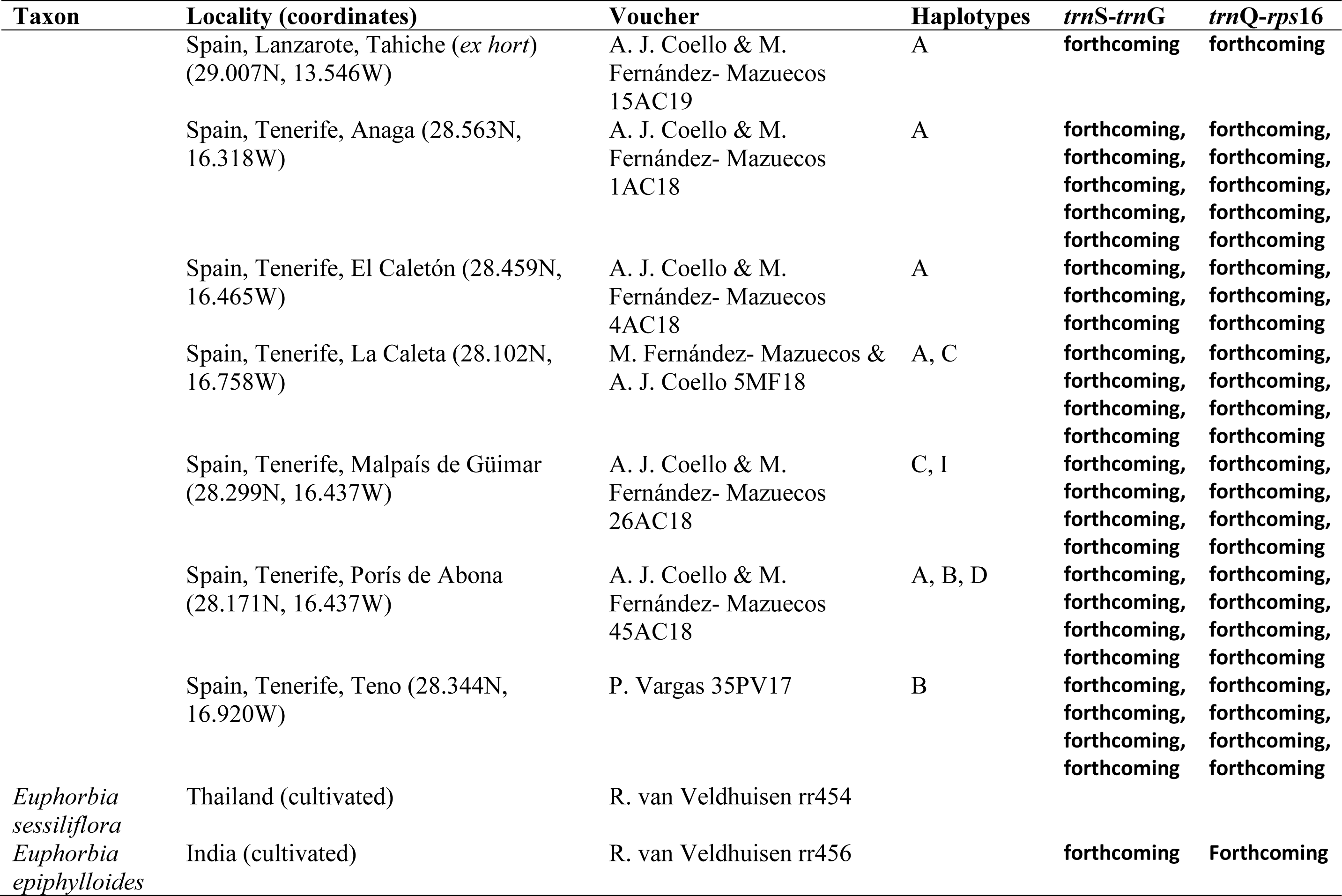
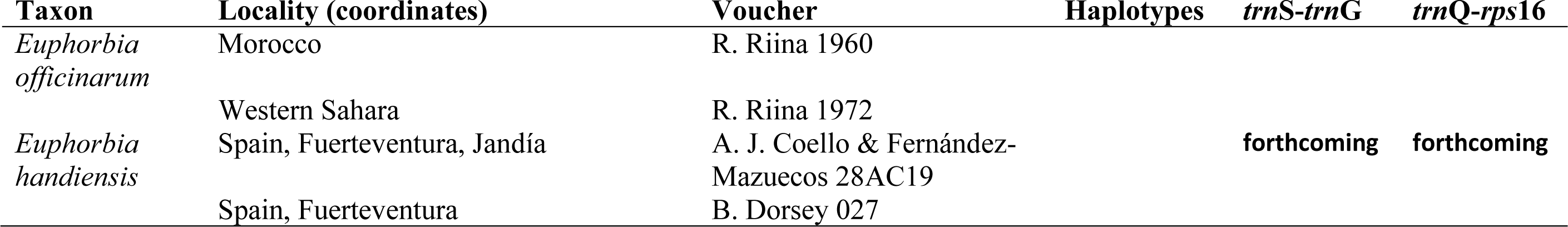
Euphorbia species used in the phylogeographic analysis of E. canariensis. Locality information (with coordinates), voucher and haplotypes detected for plastid regions are indicated.

| **Taxon** | **Locality (coordinates)** | **Voucher** | **Haplotypes** | ***trnS-trnG*** | ***trnQ-rps16*** |
| --- | --- | --- | --- | --- | --- |
| *Euphorbia abdelkuri* | Yemen (cultivated) | R. van Veldhuisen rr453 |  | forthcoming | forthcoming |
| *Euphorbia canariensis* | Spain, El Hierro, Puerto de la Estaca (27.780N, 17.906W) | A. J. Coello & M. Fernández- Mazuecos 39AC18 (MA) | G, H | forthcoming, forthcoming, forthcoming | forthcoming, forthcoming, forthcoming |
|  | Spain, El Hierro, Punta del Guincho (27.731N, 17.930W) | A. J. Coello & M. Fernández- Mazuecos 35AC18 | G | forthcoming, forthcoming, forthcoming | forthcoming, forthcoming, forthcoming |
|  | Spain, Fuerteventura, Barranco de Butihondo (28.085N, 14.325W) | A. J. Coello & M. Fernández- Mazuecos 22AC19 | A | forthcoming, forthcoming, forthcoming | forthcoming, forthcoming, forthcoming |
|  | Spain, Fuerteventura, Barranco de Vinamar (28.093N, 14.348W) | A. J. Coello & M. Fernández- Mazuecos 25AC19 | A, J | forthcoming, forthcoming, forthcoming | forthcoming, forthcoming, forthcoming |
|  | Spain, Fuerteventura, Cofete (28.096N, 14.410W) | A. J. Coello & M. Fernández- Mazuecos 27AC19 | A | forthcoming, forthcoming, forthcoming | forthcoming, forthcoming, forthcoming |
|  | Spain, Gran Canaria, Andén Verde (28.036N, 15.758W) | Y. Arjona 375YA15, 376YA15 | D | forthcoming | forthcoming |
|  | Spain, Gran Canaria, Barranco de Fataga (27.806N, 15.583W) | A. J. Coello & M. Fernández- Mazuecos 40AC19 | A | forthcoming,<br>forthcoming,<br>forthcoming,<br>forthcoming,<br>forthcoming | forthcoming,<br>forthcoming,<br>forthcoming,<br>forthcoming,<br>forthcoming |
|  | Spain, Gran Canaria, Barranco Arguineguín (27.818N, 15.664W) | Y. Arjona 393YA15, 394YA15, 395YA15, 397YA15 | C |  |  |
|  | Spain, Gran Canaria, Barranco del Cañizo (27.793N, 15.578W) | Y. Arjona 386YA15, 387YA15, 388YA15 | A | forthcoming | forthcoming |
|  | Spain, Gran Canaria, Cenobio de Valerón (28.143N, 15.607W) | A. J. Coello & M. Fernández- Mazuecos 49AC19 | A | forthcoming,<br>forthcoming,<br>forthcoming,<br>forthcoming | forthcoming,<br>forthcoming,<br>forthcoming,<br>forthcoming |
|  | Spain, Gran Canaria, La Aldea (28.009N, 15.791W) | Y. Arjona 377YA15, 380YA15, 381YA15, 382YA15 | E | forthcoming | forthcoming |
|  | Spain, Gran Canaria, La Herradura (27.996N, 15.440W) | A. J. Coello & M. Fernández- Mazuecos 35AC19 | A | forthcoming,<br>forthcoming,<br>forthcoming,<br>forthcoming | forthcoming,<br>forthcoming,<br>forthcoming,<br>forthcoming |
|  | Spain, Gran Canaria, Las Marciegas (27.997N 15.815W) | A. J. Coello & M. Fernández- Mazuecos 57AC19 | A | forthcoming,<br>forthcoming,<br>forthcoming,<br>forthcoming,<br>forthcoming | forthcoming,<br>forthcoming,<br>forthcoming,<br>forthcoming,<br>forthcoming |
|  | Spain, Gran Canaria, Mogán (27.873N, 15.731W) | Y. Arjona 401YA15, 402YA15 | A | forthcoming | forthcoming |
|  | Spain, Gran Canaria, Playa del Águila (27.778N, 15.528W) | A. J. Coello & M. Fernández- Mazuecos 44AC19 | A | forthcoming, forthcoming | forthcoming, forthcoming |
|  | Spain, La Gomera, Barranco de Arure (28.109N, 17.325W) | M. Fernández- Mazuecos & A. J. Coello 11MF18 | F | forthcoming, forthcoming, forthcoming, forthcoming | forthcoming, forthcoming, forthcoming, forthcoming |
|  | Spain, La Gomera, Punta del Joradillo (28.039N, 17.179W) | M. Fernández- Mazuecos & A.J. Coello 18MF18 | F | forthcoming, forthcoming, forthcoming | forthcoming, forthcoming, forthcoming |
|  | Spain, La Gomera, Punta Llana (28.116N, 17.112W) | A. J. Coello & M. Fernández- Mazuecos 18AC18 | F | forthcoming, forthcoming, forthcoming | forthcoming, forthcoming, forthcoming |
|  | Spain, La Palma, Fuencaliente, Los Canarios (28.495N, 17.841W) | M. Blázquez 1MBV18 | - | - | - |
|  | Spain, La Palma, Punta del Pozo (28.573N, 17.897W) | A. J. Coello & M. Fernández- Mazuecos 80AC19 | G, H | forthcoming, forthcoming, forthcoming | forthcoming, forthcoming, forthcoming |
|  | Spain, La Palma, Punta Salinas (28.742N, 17.726W) | A. J. Coello & M. Fernández- Mazuecos 67AC19 | G | forthcoming, forthcoming | forthcoming, forthcoming |
|  | Spain, La Palma, Puntallana, Caserío Lomo de los Gomereros (28.710N, 17.757W) | M. Blázquez 2MBV18 | G | forthcoming | forthcoming |
|  | Spain, La Palma, Santo Domingo (28.831N, 17.944W) | A. J. Coello & M. Fernández- Mazuecos 74AC19 | G | forthcoming, forthcoming, forthcoming | forthcoming, forthcoming, forthcoming |
|  | Spain, Lanzarote, Tahiche (*ex hort*)<br>(29.007N, 13.546W) | A. J. Coello & M.<br>Fernández- Mazuecos<br>15AC19 | A | forthcoming | forthcoming |
|  | Spain, Tenerife, Anaga (28.563N,<br>16.318W) | A. J. Coello & M.<br>Fernández- Mazuecos<br>1AC18 | A | forthcoming,<br>forthcoming,<br>forthcoming,<br>forthcoming,<br>forthcoming | forthcoming,<br>forthcoming,<br>forthcoming,<br>forthcoming,<br>forthcoming |
|  | Spain, Tenerife, El Caletón (28.459N,<br>16.465W) | A. J. Coello & M.<br>Fernández- Mazuecos<br>4AC18 | A | forthcoming,<br>forthcoming,<br>forthcoming | forthcoming,<br>forthcoming,<br>forthcoming |
|  | Spain, Tenerife, La Caleta (28.102N,<br>16.758W) | M. Fernández- Mazuecos &<br>A. J. Coello 5MF18 | A, C | forthcoming,<br>forthcoming,<br>forthcoming,<br>forthcoming | forthcoming,<br>forthcoming,<br>forthcoming,<br>forthcoming |
|  | Spain, Tenerife, Malpaís de Güimar<br>(28.299N, 16.437W) | A. J. Coello & M.<br>Fernández- Mazuecos<br>26AC18 | C, I | forthcoming,<br>forthcoming,<br>forthcoming,<br>forthcoming | forthcoming,<br>forthcoming,<br>forthcoming,<br>forthcoming |
|  | Spain, Tenerife, Porís de Abona<br>(28.171N, 16.437W) | A. J. Coello & M.<br>Fernández- Mazuecos<br>45AC18 | A, B, D | forthcoming,<br>forthcoming,<br>forthcoming,<br>forthcoming | forthcoming,<br>forthcoming,<br>forthcoming,<br>forthcoming |
|  | Spain, Tenerife, Teno (28.344N,<br>16.920W) | P. Vargas 35PV17 | B | forthcoming,<br>forthcoming,<br>forthcoming,<br>forthcoming | forthcoming,<br>forthcoming,<br>forthcoming,<br>forthcoming |
| *Euphorbia sessiliflora* | Thailand (cultivated) | R. van Veldhuisen rr454 |  |  |  |
| *Euphorbia epiphylloides* | India (cultivated) | R. van Veldhuisen rr456 |  | forthcoming | Forthcoming |
| *Euphorbia officinarum* | Morocco | R. Riina 1960 |  |  |  |
|  | Western Sahara | R. Riina 1972 |  |  |  |
| *Euphorbia handiensis* | Spain, Fuerteventura, Jandía | A. J. Coello & Fernández-Mazuecos 28AC19 |  | **forthcoming** | **forthcoming** |
|  | Spain, Fuerteventura | B. Dorsey 027 |  |  |  |

**Table S2:** Accession number of ITS sequences used in phylogenetic analysis of *Euphorbia*. For each species of *Euphorbia*, subgenus and section are indicated. New sequences generated for this study are shown in bold.

| **Species** | **Subgenus** | **Section** | **Accession number (ITS)** |
| --- | --- | --- | --- |
| *E. antso* | *Athymalus* | *Antso* | JN250112 |
| *E. aggregata* | *Athymalus* | *Athancanthae* | JN250107 |
| *E. bergeri* | *Athymalus* | *Athancanthae* | JN250119 |
| *E. bubalina* | *Athymalus* | *Athancanthae* | JN250122 |
| *E. clava* | *Athymalus* | *Athancanthae* | JN250127 |
| *E. dregeana* (1) | *Athymalus* | *Athancanthae* | JN250139 |
| *E. dregeana* (2) | *Athymalus* | *Athancanthae* | JN250140 |
| *E. fasciculata* | *Athymalus* | *Athancanthae* | JN250147 |
| *E. filiflora* | *Athymalus* | *Athancanthae* | JN250149 |
| *E. fimbriata* | *Athymalus* | *Athancanthae* | JN250150 |
| *E. flanaganii* | *Athymalus* | *Athancanthae* | JN250151 |
| *E. fusca* | *Athymalus* | *Athancanthae* | JN250155 |
| *E. gariepina* | *Athymalus* | *Athancanthae* | JN250156 |
| *E. globosa* | *Athymalus* | *Athancanthae* | JN250158 |
| *E. grantii* | *Athymalus* | *Athancanthae* | JN250162 |
| *E. heptagona* | *Athymalus* | *Athancanthae* | JN250116 |
| *E. horrida* (1) | *Athymalus* | *Athancanthae* | JN250172 |
| *E. horrida* (2) | *Athymalus* | *Athancanthae* | JN250173 |
| *E. jansenvillensis* | *Athymalus* | *Athancanthae* | JN250176 |
| *E. lignosa* | *Athymalus* | *Athancanthae* | JN250183 |
| *E. mammillaris* | *Athymalus* | *Athancanthae* | JN250188 |
| *E. namuskluftensis* | *Athymalus* | *Athancanthae* | JN250194 |
| *E. obesa* | *Athymalus* | *Athancanthae* | JN250202 |
| *E. ornithopus* | *Athymalus* | *Athancanthae* | JN250203 |
| *E. pedemontana* | *Athymalus* | *Athancanthae* | JN250208 |
| *E. polycephala* | *Athymalus* | *Athancanthae* | JN250215 |
| *E. polygona* | *Athymalus* | *Athancanthae* | JN250216 |
| *E. pseudoglobosa* | *Athymalus* | *Athancanthae* | JN250218 |
| *E. ramiglans* | *Athymalus* | *Athancanthae* | JN250224 |
| *E. schoenlandii* | *Athymalus* | *Athancanthae* | JN250234 |
| *E. balsamifera* | *Athymalus* | *Balsamis* | JN250117 |
| *E. larica* | *Athymalus* | *Balsamis* | JN207784 |
| *E. smithii* | *Athymalus* | *Lyciopsis* | JN250240 |
| *E. hadramautica* | *Athymalus* | *Pseudacalypha* | JN250166 |
| *E. longituberculosa* | *Athymalus* | *Pseudacalypha* | JN250185 |
| *E. scheffleri* | *Athymalus* | *Somalica* | JN250232 |
| *E. socrotana* | *Athymalus* | *Somalica* | JN250241 |
| *E. acerensis* | *Chamaesyce* | *Alectoroctonum* | JN250104 |
| *E. ceroderma* | *Chamaesyce* | *Alectoroctonum* | AF537389 |
| *E. cotinifolia* | *Chamaesyce* | *Alectoroctonum* | JN250130 |
| *E. dioscoreoides* | *Chamaesyce* | *Alectoroctonum* | JN250138 |
| *E. fulgens* | *Chamaesyce* | *Alectoroctonum* | JN250154 |
| *E. graminea* | *Chamaesyce* | *Alectoroctonum* | JN250159 |
| *E. graminea* 'Diamond Frost' | *Chamaesyce* | *Alectoroctonum* | JN250160 |
| *E. insulana* | *Chamaesyce* | *Alectoroctonum* | JQ750930 |
| *E. ipecacuanhae* | *Chamaesyce* | *Alectoroctonum* | JN250175 |
| *E. leucocephala* | *Chamaesyce* | *Alectoroctonum* | JN250182 |
| *E. macropus* | *Chamaesyce* | *Alectoroctonum* | AF537378 |
| *E. misera* | *Chamaesyce* | *Alectoroctonum* | JN250192 |
| *E. sciadophila* | *Chamaesyce* | *Alectoroctonum* | JN250236 |
| *E. sphaerorrhiza* | *Chamaesyce* | *Alectoroctonum* | JN250243 |
| *E. acuta* | *Chamaesyce* | *Anisophyllum* | JN250105 |
| *E. astyla* | *Chamaesyce* | *Anisophyllum* | HQ645229 |
| *E. hirta* | *Chamaesyce* | *Anisophyllum* | HQ645278 |
| *E. humifusa* | *Chamaesyce* | *Anisophyllum* | HQ645281 |
| *E. mesembryanthemifolia* | *Chamaesyce* | *Anisophyllum* | HQ645301 |
| *E. tomentulosa* | *Chamaesyce* | *Anisophyllum* | JN250250 |
| *E. ephedroides* | *Chamaesyce* | *Articulofruticosae* | JN250142 |
| *E. rhombifolia* | *Chamaesyce* | *Articulofruticosae* | JN250229 |
| *E. apparicana* | *Chamaesyce* | *Crossadenia* | JN250114 |
| *E. espinosa* | *Chamaesyce* | *Eremophyton* | AF537416 |
| *E. leistneri* | *Chamaesyce* | *Frondosae* | JN250181 |
| *E. pirottae* | *Chamaesyce* | *Frondosae* | AF537417 |
| *E. platyclada* | *Chamaesyce* | Madagascar clade | JN250213 |
| *E. cornastra* | *Chamaesyce* | *Poinsettia* | JN250129 |
| *E. cyathophora* | *Chamaesyce* | *Poinsettia* | JN250131 |
| *E. davidii* | *Chamaesyce* | *Poinsettia* | JN250136 |
| *E. eriantha* | *Chamaesyce* | *Poinsettia* | JN250144 |
| *E. exstipulata* | *Chamaesyce* | *Poinsettia* | AF537433 |
| *E. heterophylla* | *Chamaesyce* | *Poinsettia* | JN250170 |
| *E. jaliscensis* | *Chamaesyce* | *Poinsettia* | AF537442 |
| *E. pulcherrima* | *Chamaesyce* | *Poinsettia* | JN250220 |
| *E. radians* | *Chamaesyce* | *Poinsettia* | JN250223 |
| *E. scatorrhiza* | *Chamaesyce* | *Scatorrhizae* | AF537420 |
| *E. tannensis* | *Chamaesyce* | *Scatorrhizae* | JN250245 |
| *E. glanduligera* | *Chamaesyce* | *Tenellae* | JN250157 |
| *E. aphylla* | *Esula* | *Anisophyllum* | JN250113 |
| *E. bourgaeana* | *Esula* | *Anisophyllum* | JN250121 |
| *E. lateriflora* | *Esula* | *Anisophyllum* | JN250179 |
| *E. mauritanica* | *Esula* | *Anisophyllum* | JN250189 |
| *E. patula* | *Esula* | *Anisophyllum* | JN250204 |
| *E. regis-jubae* | *Esula* | *Anisophyllum* | JN250226 |
| *E. schimperi* | *Esula* | *Anisophyllum* | JN250233 |
| *E. usambarica* | *Esula* | *Anisophyllum* | KC212419 |
| *E. franchetii* | *Esula* | *Arvales* | KC212256 |
| *E. calyptrata* | *Esula* | *Calyptratae* | JN250123 |
| *E. cyparissias* | *Esula* | *Esula* | JN250133 |
| *E. ericoides* | *Esula* | *Esula* | JN250145 |
| *E. virgata* | *Esula* | *Esula* | JN250255 |
| *E. exigua* | *Esula* | *Exiguae I* | JN250146 |
| *E. rimarum* | *Esula* | *Exiguae II* | JN250230 |
| *E. guyoniana* | *Esula* | *Guyoniana* | JN250164 |
| *E. acanthothamnus* | *Esula* | *Helioscopia* | JN250103 |
| *E. akenocarpa* | *Esula* | *Helioscopia* | JN250108 |
| *E. coniosperma* | *Esula* | *Helioscopia* | KC212210 |
| *E. dimorphocaulon* | *Esula* | *Helioscopia* | JN250137 |
| *E. flavicoma* | *Esula* | *Helioscopia* | JN250152 |
| *E. helioscopia* | *Esula* | *Helioscopia* | JN250168 |
| *E. hirsuta* | *Esula* | *Helioscopia* | JN250171 |
| *E. nereidum* | *Esula* | *Helioscopia* | JN250198 |
| *E. purpurea* | *Esula* | *Helioscopia* | JN250222 |
| *E. spathulata* | *Esula* | *Helioscopia* | JN250242 |
| *E. aucheri* | *Esula* | *Herpetorrhizae* | KC212181 |
| *E. stracheyi* | *Esula* | *Holophyllum* | KC212390 |
| *E. lagascae* | *Esula* | *Lagascae* | KC212292 |
| *E. lathyris* | *Esula* | *Lathyris* | JN250180 |
| *E. myrsinites* | *Esula* | *Myrsiniteae* | JN250193 |
| *E. oxyphylla* | *Esula* | *Myrsiniteae* | JN250205 |
| *E. dendroides* | *Esula* | *Pachycladae* | JN250135 |
| *E. paralias* | *Esula* | *Paralias* | JN250207 |
| *E. segetalis* | *Esula* | *Paralias* | JN250238 |
| *E. amygdaloides* | *Esula* | *Patellares* | JN250111 |
| *E. characias* | *Esula* | *Patellares* | JN250125 |
| *E. characias* subsp.<br>*wulfenii* | *Esula* | *Patellares* | JN250126 |
| *E. kopetdaghi* | *Esula* | *Pithyusa* | JN250177 |
| *E. nicaeensis* | *Esula* | *Pithyusa* | JN250200 |
| *E. retusa* | *Esula* | *Sclerocyathium* | JN250228 |
| *E. schugnanica* | *Esula* | *Sclerocyathium* | JN250235 |
| *E. sclerocyathium* | *Esula* | *Sclerocyathium* | JN250237 |
| *E. brachycera* | *Esula* | *Tithymalus* | JN250186 |
| *E. herniariifolia* | *Esula* | *Tithymalus* | JN250169 |
| *E. peplus* (1) | *Esula* | *Tithymalus* | JN250209 |
| *E. peplus* (2) | *Esula* | *Tithymalus* | JN250210 |
| *E. abdelkuri* | *Euphorbia* | *Euphorbia* | JN250102,<br>forthcoming |
| *E. abyssinica* | *Euphorbia* | *Euphorbia* | JN207734 |
| *E. aeruginosa* | *Euphorbia* | *Euphorbia* | JN250106 |
| *E. ammak* | *Euphorbia* | *Euphorbia* | JN250110 |
| *E. canariensis* | *Euphorbia* | *Euphorbia* | **forthcoming,**<br>**forthcoming,**<br>**forthcoming,**<br>**forthcoming,**<br>**forthcoming,**<br>**forthcoming** |
| *E. drupifera* | *Euphorbia* | *Euphorbia* | JN250141 |
| *E. epiphylloides* | *Euphorbia* | *Euphorbia* | JN250143,<br>**forthcoming** |
| *E. grandicornis* | *Euphorbia* | *Euphorbia* | JN250161 |
| *E. griseola* | *Euphorbia* | *Euphorbia* | JN250163 |
| *E. gymnocalycoides* | *Euphorbia* | *Euphorbia* | JN250165 |
| *E. handiensis* | *Euphorbia* | *Euphorbia* | **forthcoming,**<br>**forthcoming** |
| *E. ingens* | *Euphorbia* | *Euphorbia* | JN207781 |
| *E. lactea* | *Euphorbia* | *Euphorbia* | JN250178 |
| *E. micracantha* | *Euphorbia* | *Euphorbia* | JN250190 |
| *E. neriifolia* | *Euphorbia* | *Euphorbia* | JN250199 |
| *E. nyassae* | *Euphorbia* | *Euphorbia* | JN250201 |
| *E. officinarum* | *Euphorbia* | *Euphorbia* | **forthcoming,**<br>**forthcoming** |
| *E. piscidermis* | *Euphorbia* | *Euphorbia* | JN250212 |
| *E. ramipressa* | *Euphorbia* | *Euphorbia* | JN250225 |
| *E. resinifera* | *Euphorbia* | *Euphorbia* | JN250227 |
| *E. robecchii* | *Euphorbia* | *Euphorbia* | JN250231 |
| *E. sessiliflora* | *Euphorbia* | *Euphorbia* | **forthcoming** |
| *E. teke* | *Euphorbia* | *Euphorbia* | JN250246 |
| *E. turbiniformis* | *Euphorbia* | *Euphorbia* | JN250251 |
| *E. unispina* | *Euphorbia* | *Euphorbia* | JN250253 |
| *E. venenifica* | *Euphorbia* | *Euphorbia* | JN250254 |
| *E. alluaudii* | *Euphorbia* | *Goniostema* + *Deuterococalli* + *Denisophorbia* | JN250109 |
| *E. beharensis* var. *guillemetii* | *Euphorbia* | *Goniostema* + *Deuterococalli* + *Denisophorbia* | JN250118 |
| *E. cylindrifolia* | *Euphorbia* | *Goniostema* + *Deuterococalli* + *Denisophorbia* | JN250132 |
| *E. decaryi* | *Euphorbia* | *Goniostema* + *Deuterococalli* + *Denisophorbia* | JN250134 |
| *E. francosisii* | *Euphorbia* | *Goniostema* + *Deuterococalli* + *Denisophorbia* | JN250153 |
| *E. hedyotoides* | *Euphorbia* | *Goniostema* + *Deuterococalli* + *Denisophorbia* | JN250167 |
| *E. iharanae* | *Euphorbia* | *Goniostema* + *Deuterococalli* + *Denisophorbia* | JN250174 |
| *E. milii* | *Euphorbia* | *Goniostema* + *Deuterococalli* + *Denisophorbia* | JN250191 |
| *E. neohumbertii* | *Euphorbia* | *Goniostema* + *Deuterococalli* + *Denisophorbia* | JN250196 |
| *E. neoarborescens* | *Euphorbia* | *Monadenium* | JN250195 |
| *E. neospinescens* | *Euphorbia* | *Monadenium* | JN250197 |
| *E. umbellata* | *Euphorbia* | *Monadenium* | AF537469 |
| *E. boophthona* | *Euphorbia* | New World Clade + *Pacifcae* | JN250120 |
| *E. calyculata* | *Euphorbia* | New World Clade + *Pacifcae* | KJ888151 |
| *E. comosa* | *Euphorbia* | New World Clade + *Pacifcae* | JN250128 |
| *E. germainii* | *Euphorbia* | New World Clade + *Pacifcae* | AF537499 |
| *E. lactiflua* | *Euphorbia* | New World Clade + *Pacifcae* | AF537528 |
| *E. lomelii* | *Euphorbia* | New World Clade + *Pacifcae* | JN250184 |
| *E. plumerioides* | *Euphorbia* | New World Clade + *Pacifcae* | JN250214 |
| *E. portulacoides* subsp. *collina* | *Euphorbia* | New World Clade + *Pacifcae* | JN250217 |
| *E. pteroneura* | *Euphorbia* | New World Clade + *Pacifcae* | JN250219 |
| *E. punicea* | *Euphorbia* | New World Clade + *Pacifcae* | JN250221 |
| *E. sinclairiana* | *Euphorbia* | New World Clade + *Pacifcae* | AF537495 |
| *E. sipolisii* | *Euphorbia* | New World Clade + *Pacifcae* | JN250239 |
| *E. tanquahuete* | *Euphorbia* | New World Clade + *Pacifcae* | AF537525 |
| *E. tithymaloides* | *Euphorbia* | New World Clade + *Pacifcae* | JN250249 |
| *E. umbelliformis* | *Euphorbia* | New World Clade + *Pacifcae* | JN250252 |
| *E. weberbaueri* | *Euphorbia* | New World Clade + *Pacifcae* | JN250256 |
| *E. pachysantha* | *Euphorbia* | *Pachysanthae* | JN250206 |
| *E. brunellii* | *Euphorbia* | *Rubellae* | AF537486 |
| *E. rubella* | *Euphorbia* | *Rubellae* | AF537487 |
| *E. arbuscula* | *Euphorbia* | *Tirucalli + Pervilleanae* | JN250115 |
| *E. fiherenensis* | *Euphorbia* | *Tirucalli + Pervilleanae* | JN250148 |
| *E. pervilleana* | *Euphorbia* | *Tirucalli + Pervilleanae* | JN250211 |
| *E. stenoclada* | *Euphorbia* | *Tirucalli + Pervilleanae* | JN250244 |
| *E. tirucalli* (1) | *Euphorbia* | *Tirucalli + Pervilleanae* | JN250247 |
| *E. tirucalli* (2) | *Euphorbia* | *Tirucalli + Pervilleanae* | JN250248 |
| *Anthostema senegalense* |  |  | JN250088 |
| *Bonania microphylla* |  |  | JN250089 |
| *Calycopeplus casuarinoides* |  |  | AF537580 |
| *Colliguaja integerrima* |  |  | JN250090 |
| *Dichostemma glaucescens* |  |  | JN250091 |
| *Gymnanthes albicans* |  |  | JN250092 |
| *Homalanthus nutans* |  |  | JN250093 |
| *Hura crepitans* |  |  | JN250094 |
| *Mabea* sp. |  |  | JN250095 |
| *Maprounea guianensis* |  |  | JN250096 |
| *Microstachys chamaelea* |  |  | JN250097 |
| *Nealchornea yapurensis* |  |  | JN250098 |
| *Neoguillauminia cleopatra* |  |  | JN250099 |
| *Senefelderopsis croizatii* |  |  | JN250100 |
| *Stillingia sylvatica* subsp. *tenuis* |  |  | JN250101 |

**Table S3:** Final set of E. canariensis occurrences used for SDM analysis.

| **Island** | **Latitude** | **Longitude** | **Island** | **Latitude** | **Longitude** |
| --- | --- | --- | --- | --- | --- |
| El Hierro | 27.77984 | -17.90645 | La Palma | 28.74249 | -17.72608 |
| El Hierro | 27.68851051 | -17.96125023 | La Palma | 28.83092 | -17.94396 |
| El Hierro | 27.83851155 | -17.9312516 | La Palma | 28.639985 | -17.75842 |
| Fuerteventura | 28.08461 | -14.3253 | La Palma | 28.840575 | -17.78821 |
| Fuerteventura | 28.0964 | -14.40969 | La Palma | 28.654893 | -17.94662 |
| Gran Canaria | 28.03553894 | -15.7579237 | La Palma | 28.835178 | -17.88413 |
| Gran Canaria | 27.80637 | -15.58282 | La Palma | 28.753718 | -18.00421 |
| Gran Canaria | 27.81756294 | -15.6642949 | Tenerife | 28.56301 | -16.31792 |
| Gran Canaria | 28.14286 | -15.60717 | Tenerife | 28.45859 | -16.46548 |
| Gran Canaria | 27.99632 | -15.44031 | Tenerife | 28.10211 | -16.75767 |
| Gran Canaria | 27.99694 | -15.81488 | Tenerife | 28.29871 | -16.37118 |
| Gran Canaria | 27.87292776 | -15.7307223 | Tenerife | 28.17112 | -16.43661 |
| Gran Canaria | 27.77808 | -15.52824 | Tenerife | 28.343889 | -16.91972 |
| Gran Canaria | 27.9437 | -15.4309 | Tenerife | 28.56911 | -16.25391 |
| Gran Canaria | 28.069237 | -15.462275 | Tenerife | 28.250997 | -16.44106 |
| Gran Canaria | 28.08643 | -15.69589 | Tenerife | 28.5323 | -16.1587 |
| Gran Canaria | 27.784606 | -15.718496 | Tenerife | 28.028642 | -16.70458 |
| Gran Canaria | 27.922775 | -15.81061 | Tenerife | 28.401027 | -16.58177 |
| La Gomera | 28.10924 | -17.32529 | Tenerife | 28.117373 | -16.50807 |
| La Gomera | 28.03922 | -17.17908 | Tenerife | 28.294594 | -16.84741 |
| La Gomera | 28.11556 | -17.11221 | Tenerife | 28.032724 | -16.59973 |
| La Gomera | 28.175226 | -17.177349 | Tenerife | 28.113777 | -16.68097 |
| La Gomera | 28.0585193 | -17.2312446 | Tenerife | 28.244393 | -16.83393 |
| La Gomera | 28.11852019 | -17.1712445 | Tenerife | 28.378809 | -16.81564 |
| La Gomera | 28.05851883 | -17.2812453 | Tenerife | 28.383751 | -16.373 |
| La Palma | 28.495466 | -17.840561 | Tenerife | 28.518723 | -16.22791 |
| La Palma | 28.57313 | -17.8974 |  |  |  |

**Table S4:** Climatic variables considered for SDM analysis of *E. canariensis* before variable selection.

| **Variable** | **Abbreviation** |
| --- | --- |
| Annual mean temperature | bio 1 |
| Mean diurnal range (mean of monthly (max temp – min temp)) | bio 2 |
| Isothermality (bio 2/bio 7) ($\times 100$) | bio 3 |
| Temperature seasonality (standard deviation $\times 100$) | bio 4 |
| Max temperature of warmest month | bio 5 |
| Min temperature of coldest month | bio 6 |
| Temperature annual range (bio 5 – bio 6) | bio 7 |
| Mean temperature of wettest quarter | bio 8 |
| Mean temperature of driest quarter | bio 9 |
| Mean temperature of warmest quarter | bio 10 |
| Mean temperature of coldest quarter | bio 11 |
| Annual precipitation | bio 12 |
| Precipitation of wettest month | bio 13 |
| Precipitation of driest month | bio 14 |
| Precipitation seasonality (coefficient of variation) | bio 15 |
| Precipitation of wettest quarter | bio 16 |
| Precipitation of driest quarter | bio 17 |
| Precipitation of warmest quarter | bio 18 |
| Precipitation of coldest quarter | bio 19 |
| Bulk density (fine earth) | eda 1 |
| Volumetric percentage of coarse fragments ($>2$ mm) | eda 2 |
| Weight percentage of clay particles ($<0.0002$ mm) | eda 3 |
| Weight percentage of silt particles ($0.0002$–$0.05$ mm) | eda 4 |
| Weight percentage of sand particles ($0.05$–$2$ mm) | eda 5 |
| Cation exchange capacity of soil | eda 6 |
| Soil organic carbon content | eda 7 |
| Soil organic carbon stock | eda 8 |
| pH index measured in water solution | eda 9 |
| pH index measured in KCl solution | eda 10 |

**Figure S1:**
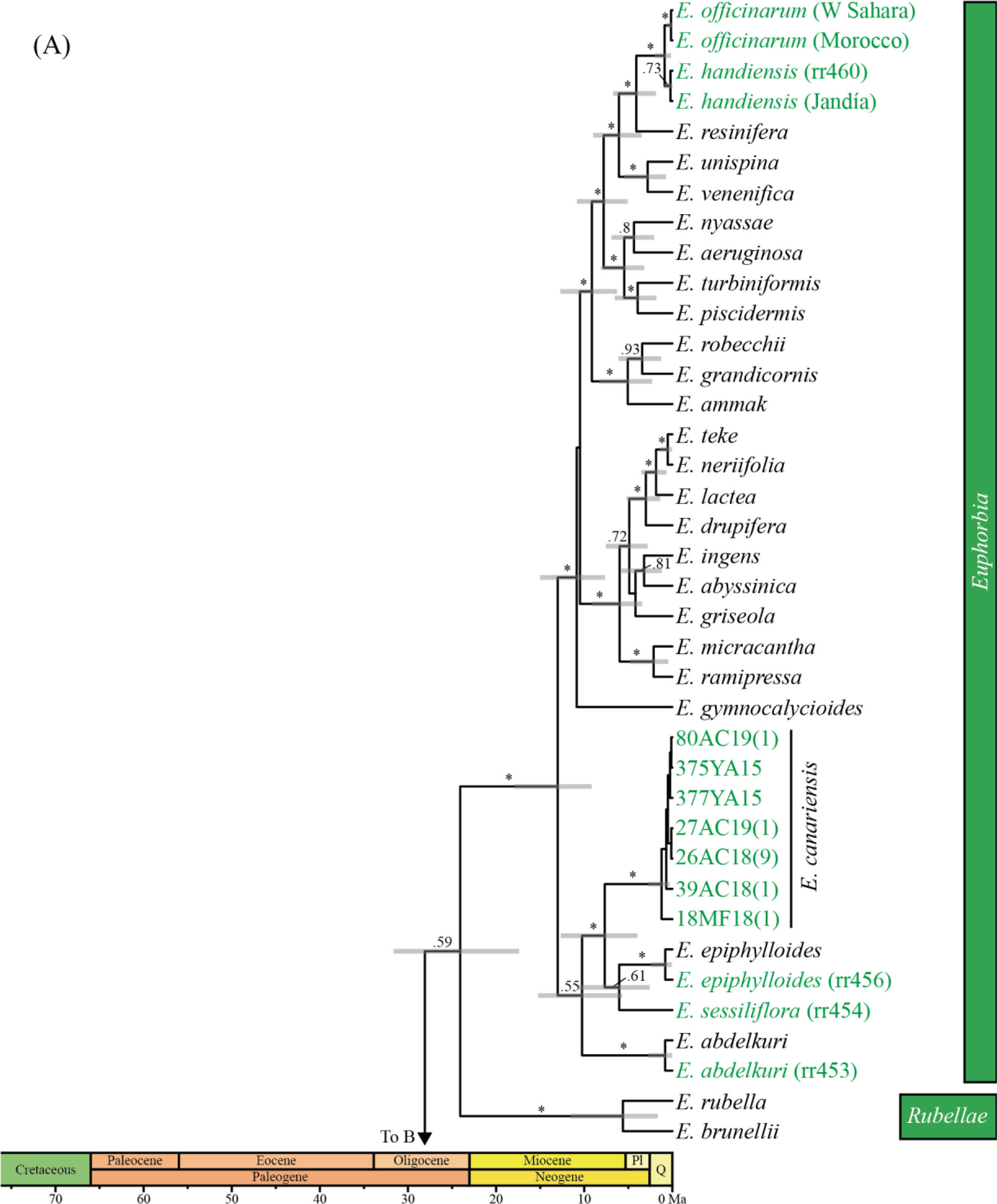
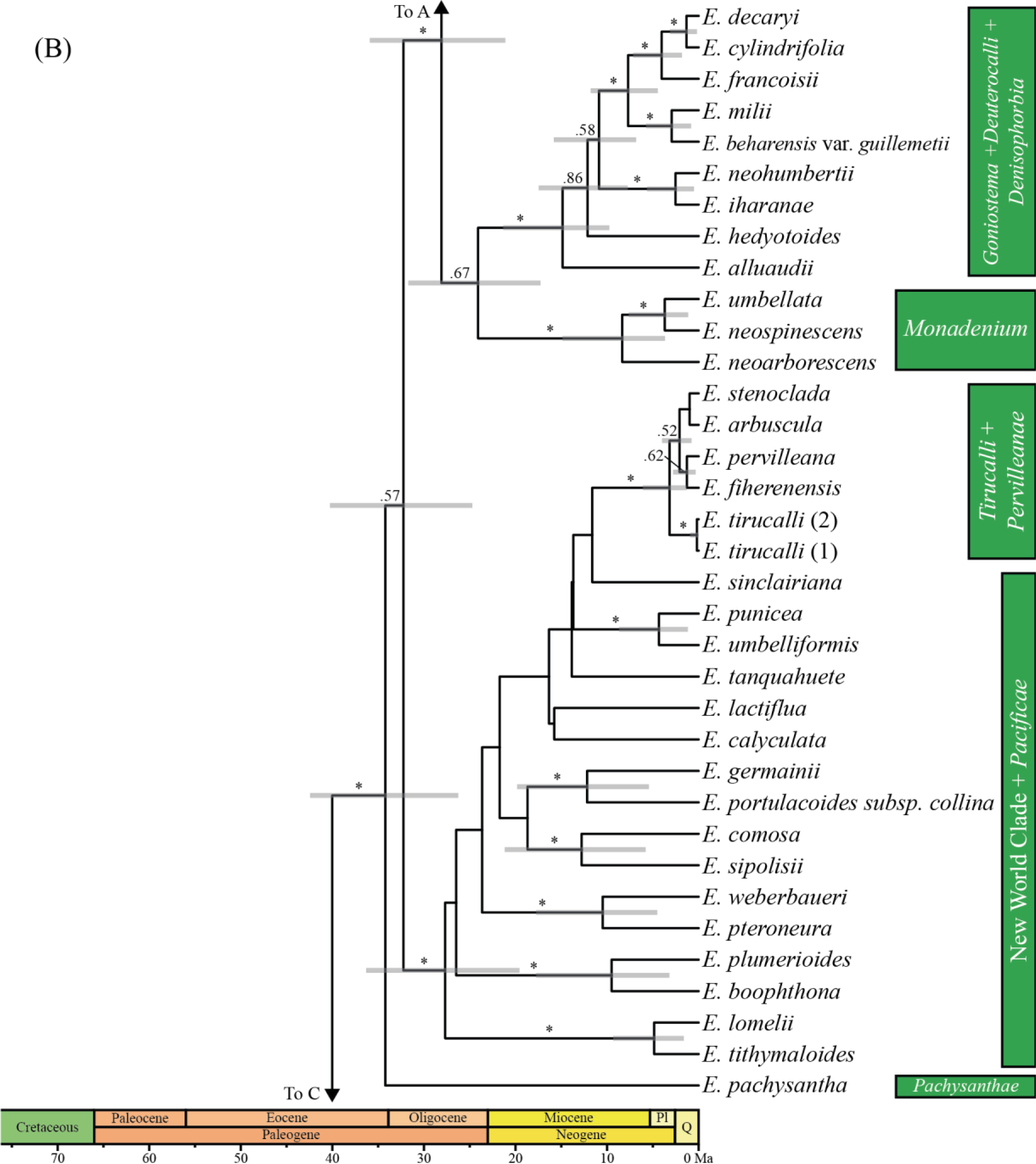
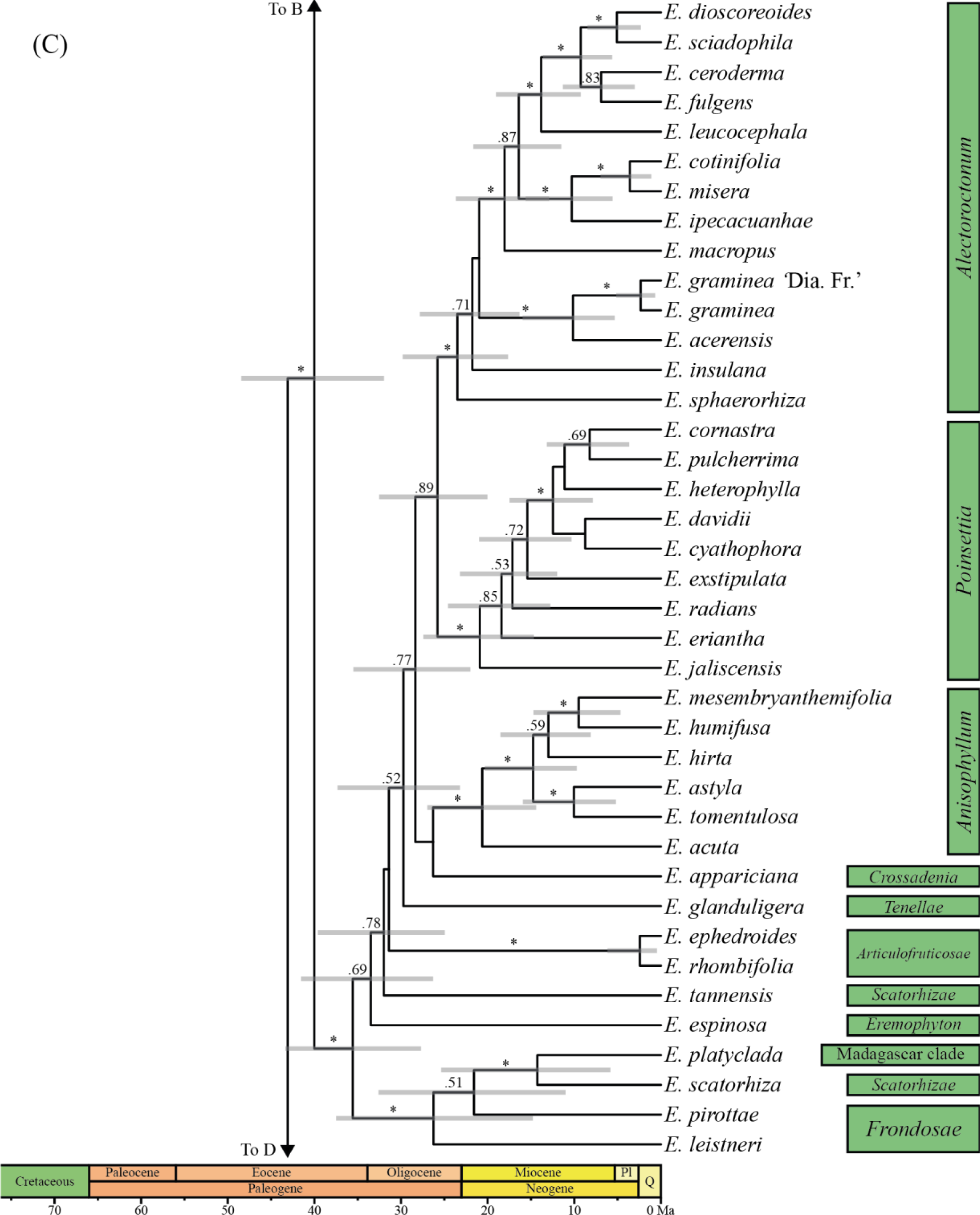
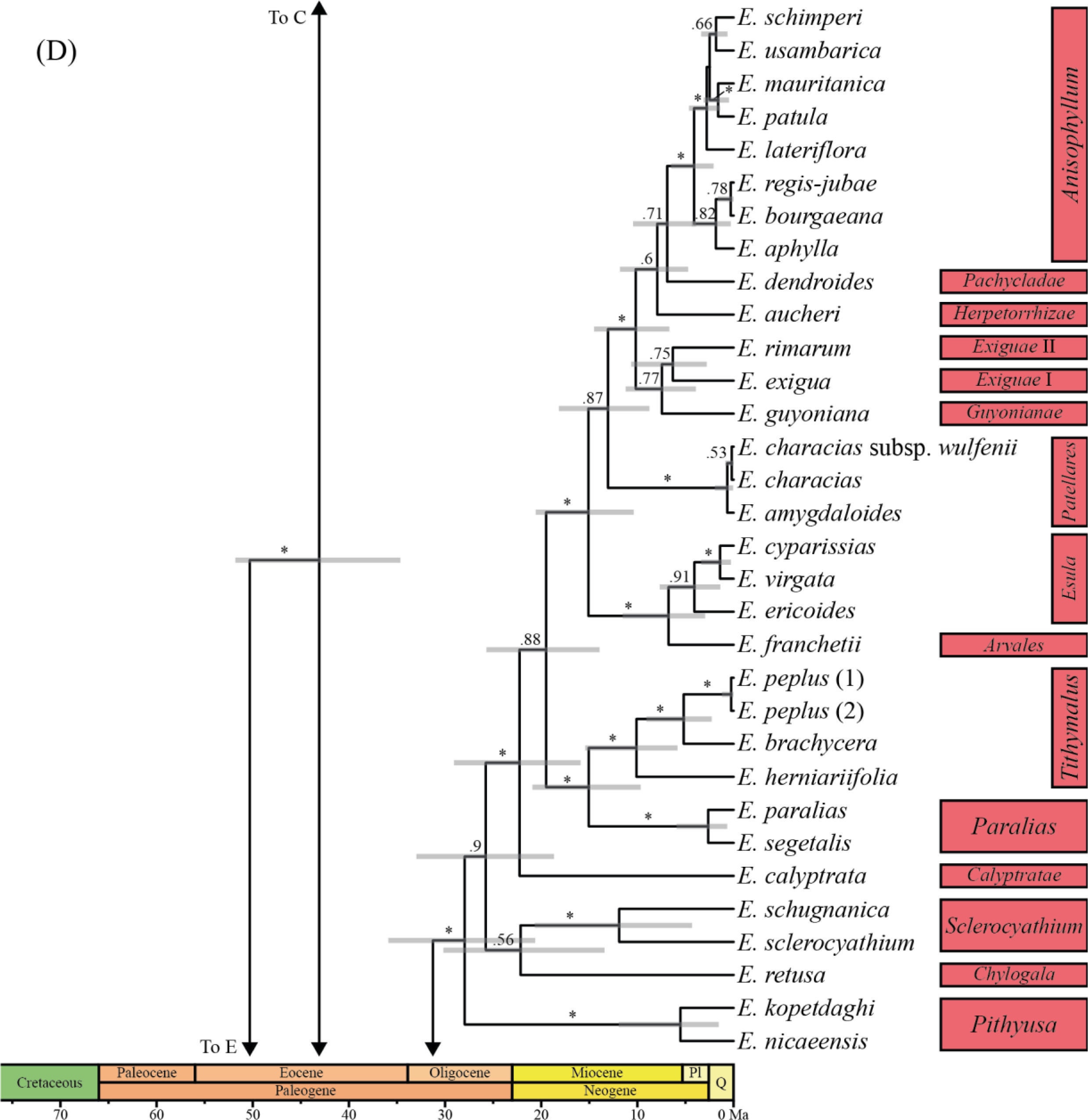
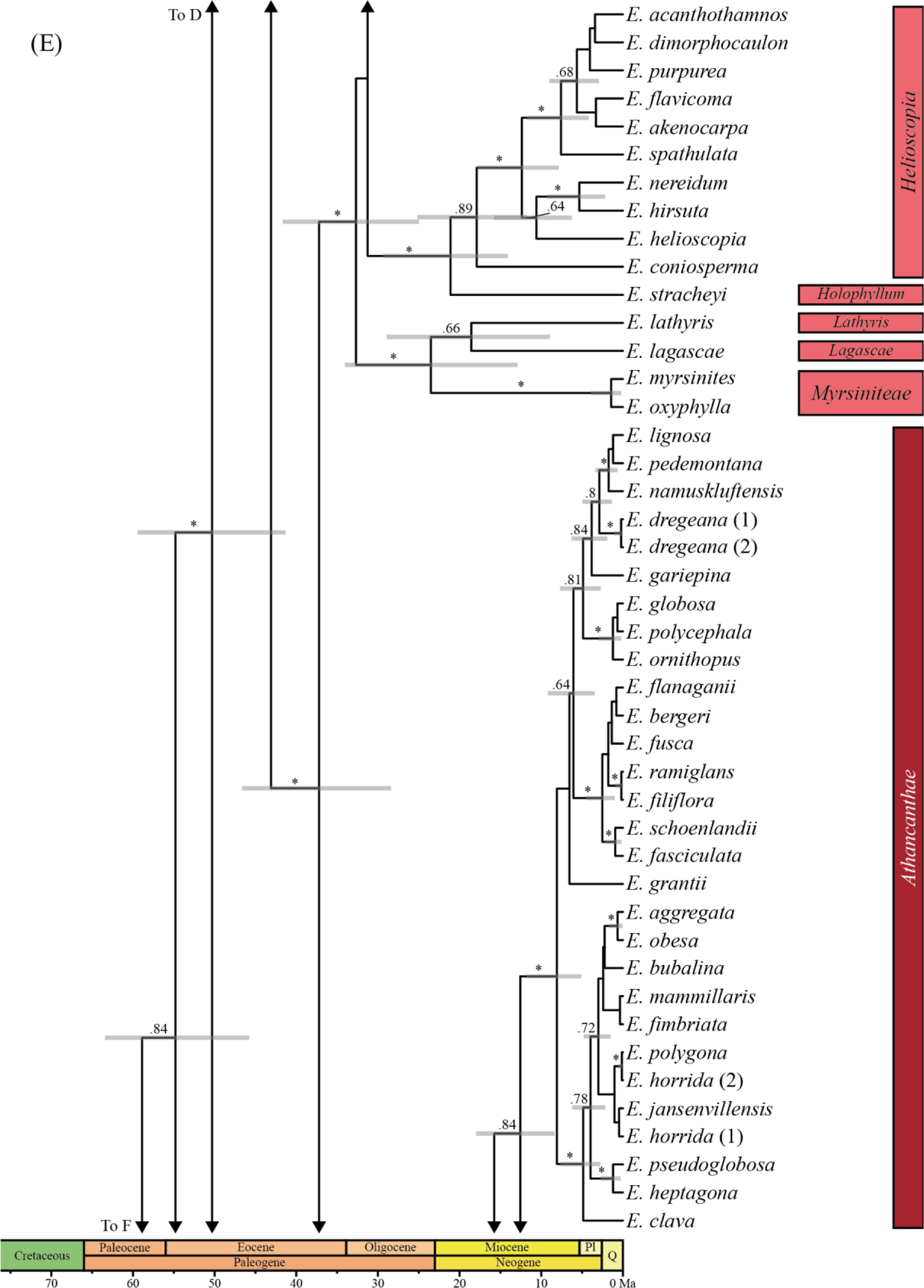
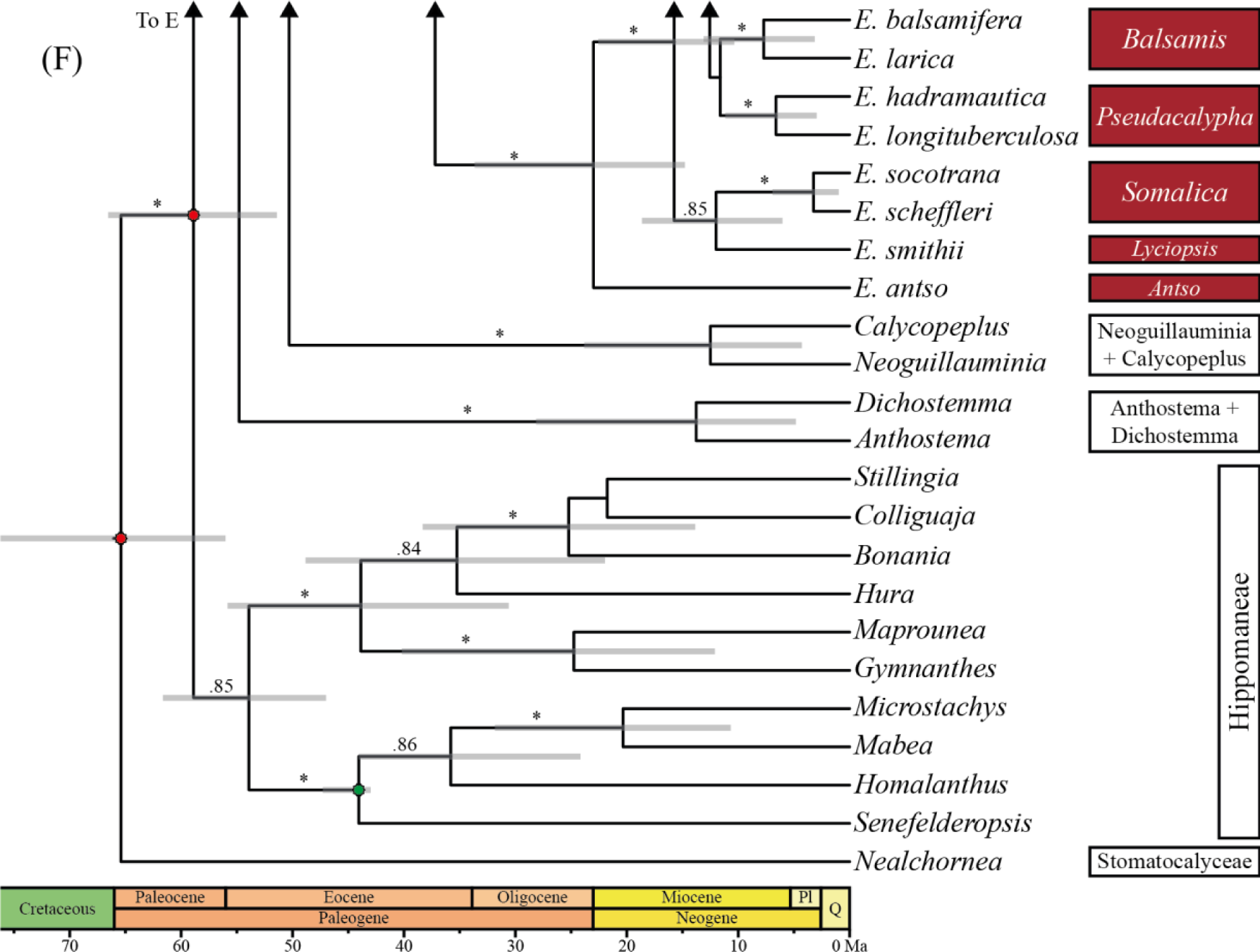
Complete Bayesian chronogram of *Euphorbia*. Species are grouped according to their section following Horn et al. (2014). Terminals in green are those sequenced for this study, while all others were obtained from a previous phylogeny (Horn et al., 2014). Section labels are coloured according to the four subgenera of *Euphorbia*: *Euphorbia* in dark green, *Chamaesyce* in light green, *Esula* in light red, and *Athymalus* in dark red. Outgroup labels are shown in white. Calibration points are indicated as stars (red for secondary calibrations and green for the fossil calibration; see text for further details). Numbers above branches indicate posterior probabilities (PP) when PP > 0.5, and asterisks (*) indicate PP > 0.95. Node bars indicates the 95% highest posterior density intervals for ages when PP > 0.5.

**Alignment S1:** Alignment of Euphorbia phylogeny analysis.

**Alignment S2:** Alignment of *Euphorbia* haplotypes for phylogeographic analysis.

## Data availability

Data is available in Genbank and in the online version of this article. Further information is available on request.

## Notes

### Competing Interest Statement

The authors have declared no competing interest.

